# Uncertainty-aware benchmarking reveals ambiguous transcripts in mRNA–lncRNA classification

**DOI:** 10.64898/2026.04.14.718168

**Authors:** Daniel Garcia-Ruano, Mikaël Georges, Saswat K. Mohanty, Rahma Baaziz, Kateryna D. Makova, Macha Nikolski, Domitille Chalopin

## Abstract

**Background:** Long non-coding RNAs (lncRNAs) have gained significant attention in recent years, yet distinguishing them from protein-coding transcripts remains challenging. Indeed, many lncRNAs share mRNA-like processing and existing sequence-derived signals do not fully capture the coding/non-coding boundary. Recent GENCODE annotation efforts revealed tens of thousands of novel lncRNA sequences as well as the reclassification of some lncRNAs into the protein-coding class, highlighting the need to better characterize transcript features associated with classification uncertainty and errors.

**Results:** We performed uncertainty-aware benchmarking by retraining and evaluating eight transcript classifiers under a controlled protocol on a label-stable GENCODE v46–v47 subset. Beyond conventional model evaluation metrics, we quantified inter-tool agreement and entropy-based uncertainty to stratify transcripts into consensus, discordant, and consensus-error groups. To expand standard sequence and ORF-derived signals, we incorporated repeat-derived features from mature transcripts and non-B DNA motif features across gene bodies. Although aggregate performance was high, ∼45% of transcripts showed inter-tool discordance, particularly among lncRNAs. Feature analyses linked low-uncertainty predictions to strong coding-like signals, whereas high-uncertainty profiles exhibited mixed signatures. Alongside classical predictors in global importance analyses, repeat-derived features appear as main contributors.

**Conclusions:** By combining controlled benchmarking with transcript-level agreement and uncertainty stratification, together with extended feature profiling, we identified patterns associated with classifier disagreement and misclassification. This novel framework provides practical guidance for interpreting predictions, motivating the development of more robust coding/non-coding classifiers, while also shedding light on the sequence properties that distinguish lncRNA sequences.

## Background

Long non-coding RNAs (lncRNAs) constitute a large and heterogeneous class of non-coding RNAs longer than 200 nucleotides, encompassing more than 35,000 annotated human loci in the latest GENCODE update [1,2]. Recognized as regulators of gene expression, they exert their functions through interactions with DNA, RNA and proteins, influencing chromatin organization, RNA splicing, stability and translation [3,4]. With their ability to regulate gene expression at multiple levels, lncRNAs are central players in many biological processes, such as X-chromosome inactivation or cell differentiation. However, their functional diversity, high isoform variability, and numerous similarities with mRNAs such as splicing, capping, and polyadenylation, make them difficult to distinguish from protein-coding transcripts purely from transcript sequence. Additionally, the presence of antisense transcripts and sometimes their processing from intronic or repetitive sequences further blurs the boundary between protein-coding and non-coding genes.

Distinguishing these two types of transcripts has long challenged computational genomics. Early approaches such as Fickett’s TESTCODE algorithm [5] detected coding regions *ab initio*. With the recognition of the pervasive transcription of non-coding genomic regions, tools such as PhyloCSF [6], CPC [7] and CPAT [8] were developed to classify transcripts using evolutionary conservation and sequence composition. Over the past decade, machine learning methods have incorporated broader feature sets, including k-mer frequencies, RNA secondary structure, and predicted protein properties; and, more recently, deep learning models have been trained on raw transcript sequences. Although these methods can achieve high accuracy on standard within-species benchmarks, performance can degrade substantially under class imbalance, cross-species transfer, and challenging transcript profiles (e.g., extreme length or complex isoform structure) [9]. Recently, lncRNA-BERT leveraged transformer architectures and reported strong performance on several commonly used benchmarks [10]. However, important limitations remain, both in terms of data complexity and model limitations. This includes annotation updates, dataset shifts, sequence redundancy, context-length constraints for long transcripts, cross-species model generalization and limited biological interpretability.

As these classifiers proliferated, benchmarking studies have increasingly moved beyond analyzing accuracy to exploring their biological interpretability. For example, Klapproth et al. [11] argued that most classifiers detect the absence of coding signals rather than lncRNA-specific properties. Others reported systematic biases, such as performance drops for sequences outside typical length ranges of 1-10 kb [11], and consistent misclassification of particular transcript subsets across dozens of models [12]. While these studies provided valuable insights, analyses often focused on a narrow set of transcript properties (e.g., length). Moreover, many large-scale benchmarks provide limited control over dataset composition, which can introduce data leakage via redundancy, near-duplicate isoforms, or overlapping annotations between training and evaluation sets. As a result, key questions remain: to what extent do state-of-the-art tools produce concordant predictions on the same transcripts? Do “difficult” transcripts share distinctive and interpretable properties? And can features beyond sequence-derived statistics help explain disagreement and improve classification?

In this study, we address these limitations by (1) retraining eight representative coding-potential classifiers under a controlled protocol, (2) quantifying prediction confidence together with inter-model agreement to stratify transcripts into operational “difficulty groups”, and (3) analyzing the features associated with confident, correct and discordant predictions (Fig. 1). Beyond the raw transcript sequences, secondary structure and predicted protein characteristics, we incorporate additional feature sets, including non-B DNA motifs, predicted from the genomic sequence and known to modulate transcription [13–16]; and repetitive element content from mature transcripts, potentially influencing the functional properties of the RNA itself. This framework supports a more robust and reproducible comparison of tools, provides practical guidance on how to interpret model disagreements, and highlights transcript types that remain challenging, not only suggesting directions for future model development but also providing insights on lncRNA complexity.

**Figure 1.**
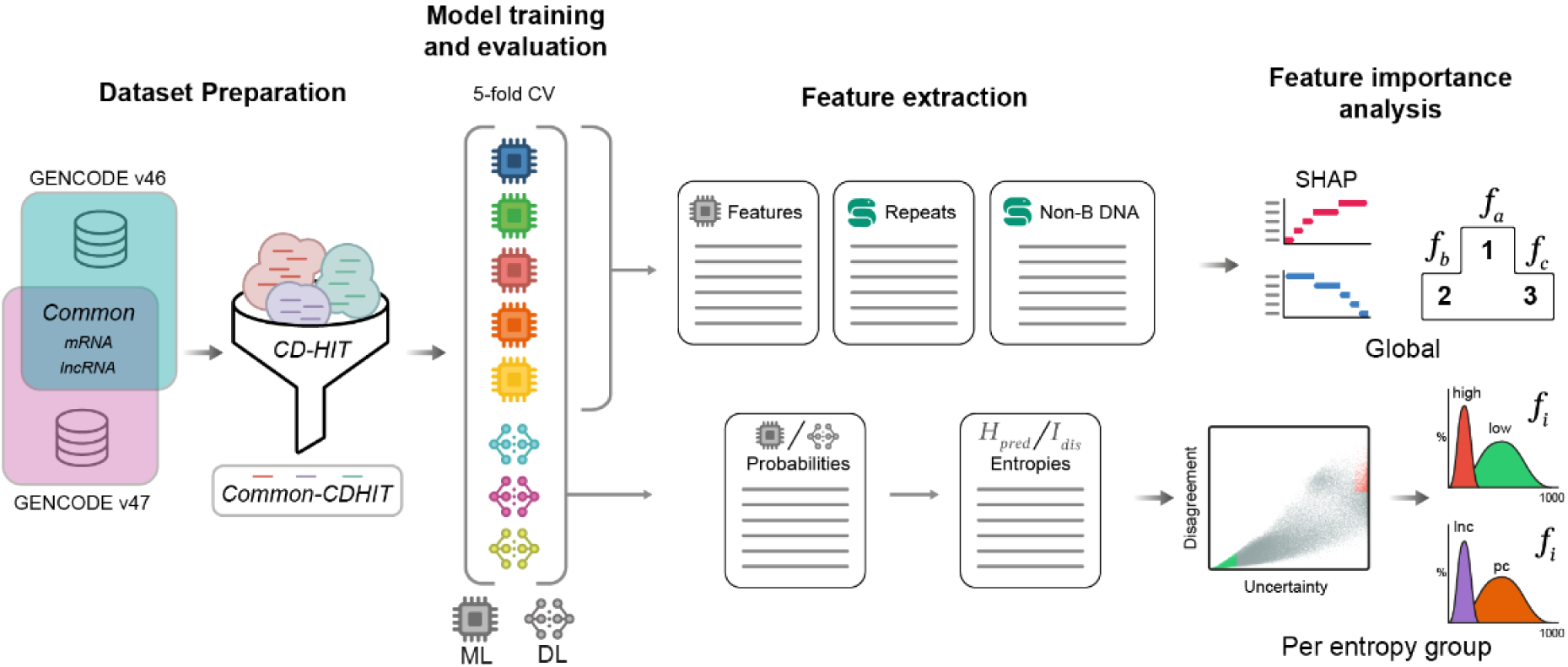
Overview of the benchmark approach: Identifying Classification Failures via Entropy and Atypical Structures.

## Results

### Construction of a high-quality, non-redundant benchmarking dataset and extraction of genomic features

Fair comparison of coding/non-coding classifiers requires evaluation on identical train/test sets and explicit control over train-test leakage due to sequence redundancy (e.g., highly similar isoforms). However, most available models are trained on unpublished or outdated datasets, either making it difficult to retrain them under identical conditions and to quantify leakage and dataset shift.

To mitigate these issues in our study, we constructed a benchmarking dataset with 3 design principles: (1) annotation stability and curation, to reduce label noise; (2) reduced sequence redundancy, to limit near-duplicate information leakage; and (3) balanced training classes (i.e., equal number of protein-coding and lncRNA transcripts), to avoid model learning a trivial majority class detection rule.

We used Human GENCODE v47 as the curated reference annotation for human transcripts. In a recent publication of the GENCODE consortium, Capture Long-Read Sequencing (CLS) combined with the updated TAGADA pipeline added 140,268 novel lncRNA transcripts (and 17,931 novel loci) in v47 compared to v46, roughly doubling the lncRNA catalog (Kaur et al., 2024; Mudge et al., 2025) (Fig. 2A, B). Because GENCODE labels are refined over time, transcripts annotated as lncRNA in one release can later be reassigned as protein-coding (or vice versa) as evidence accumulates. To reduce label instability, we defined a “common” subset comprising transcripts that (1) are present in both v46 and v47, (2) are labeled as either protein-coding or lncRNA, and (3) retain the same class label across releases (Table 1, Methods). This yields a conservative benchmark set focused on stable labels rather than maximal catalogue size.

**Figure 2.**
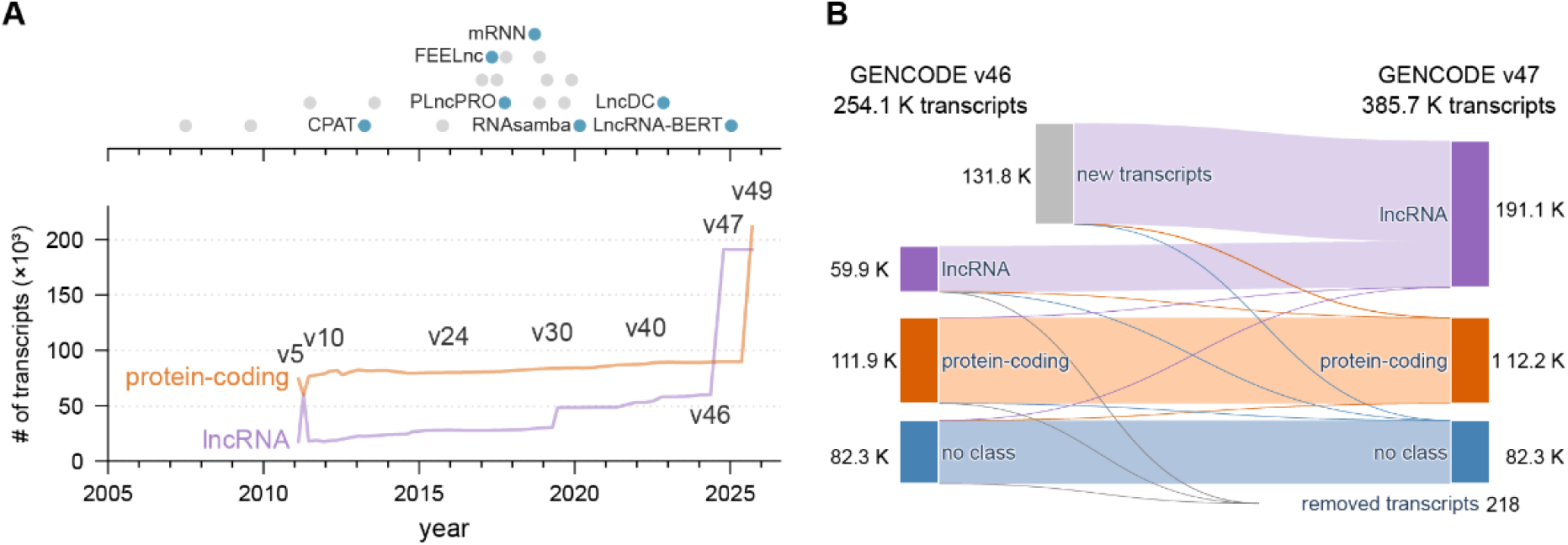
Evolution of transcript annotations and classification tools. (A) Timeline of transcript classification tools by publication date, with blue highlighting those benchmarked in this study (top). Number of human protein-coding and lncRNA transcripts annotated across all GENCODE database versions, with selected version numbers labelled for reference (bottom). Tools displayed in A are detailed in Table S1. (B) Sankey plot showing changes in coding class annotation (lncRNA, protein-coding and no class) between versions v46 and v47 of the GENCODE database, including transcripts added or removed in v47. “No class” transcripts are those which do not fit in either of the protein-coding or lncRNA groups, such as short non-coding RNAs. Link widths are proportional to transcript numbers, illustrating the stability and transitions between versions. Data used to construct B are detailed in Table S2.

**Table 1.**
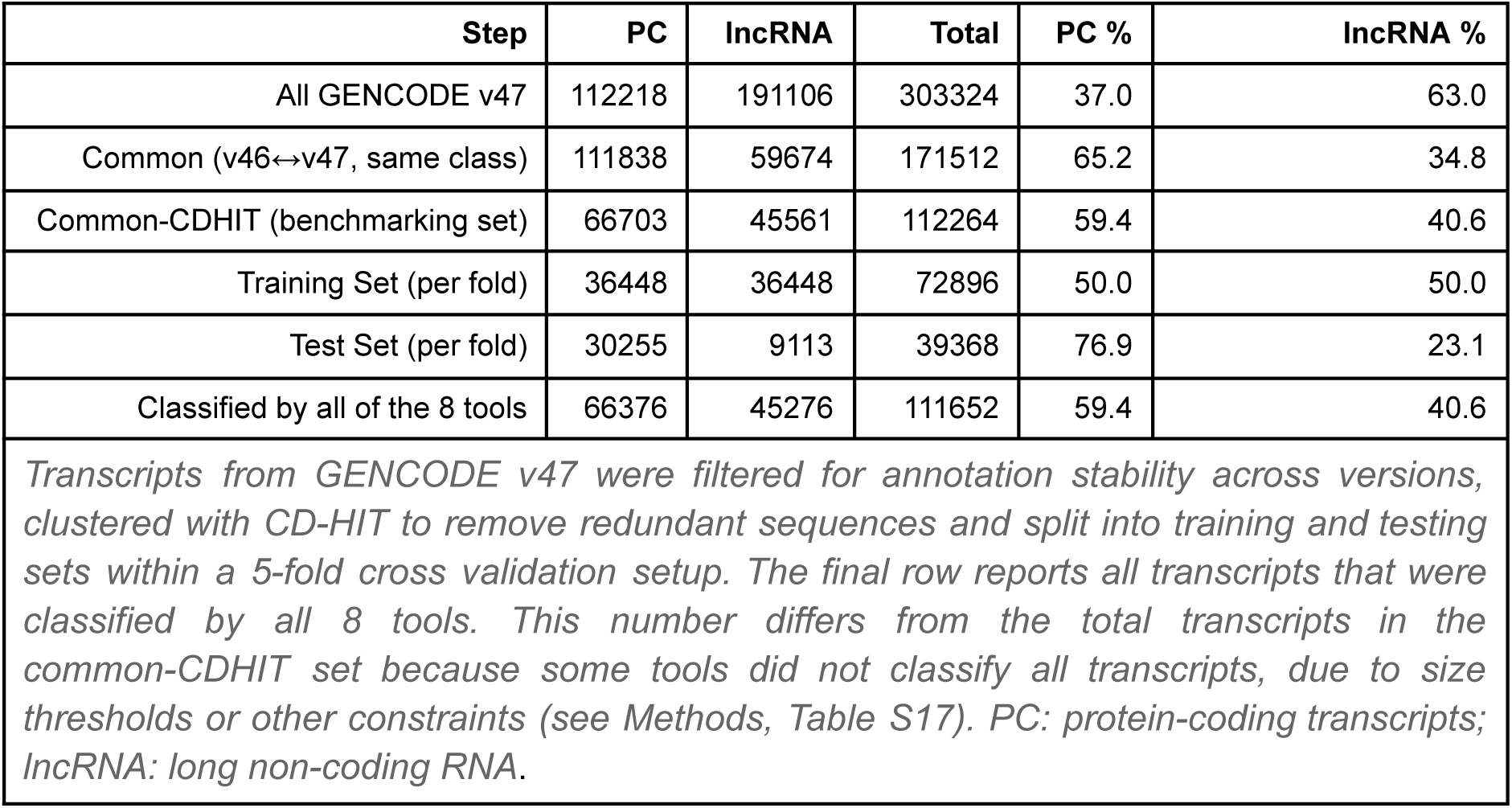
Protein-coding and lncRNA transcript counts at each processing step.

|  |  |  |  |  |  |
| --- | --- | --- | --- | --- | --- |
| Step | PC | lncRNA | Total | PC % | lncRNA % |
| All GENCODE v47 | 112218 | 191106 | 303324 | 37.0 | 63.0 |
| Common (v46↔v47, same class) | 111838 | 59674 | 171512 | 65.2 | 34.8 |
| Common-CDHIT (benchmarking set) | 66703 | 45561 | 112264 | 59.4 | 40.6 |
| Training Set (per fold) | 36448 | 36448 | 72896 | 50.0 | 50.0 |
| Test Set (per fold) | 30255 | 9113 | 39368 | 76.9 | 23.1 |
| Classified by all of the 8 tools | 66376 | 45276 | 111652 | 59.4 | 40.6 |

To reduce sequence redundancy and minimize near-duplicate leakage, we clustered transcripts in the “common” set using CD-HIT [17], as is common practice in coding-potential tool development (e.g., CPC2, lncDC, lncRNA-BERT). Shortly, CD-HIT groups sequences into the same cluster based on a 90% similarity with the representative sequence. 90% identity threshold over the shortest transcript We selected the longest sequence per cluster as representative, ensuring maximal retention of exon content while avoiding redundant training examples. Our final dataset, named “**common-CDHIT**” contains 45,561 lncRNAs and 66,703 mRNAs, that were later stratified in five cross-validation fold subsets, in which the training set was class-balanced to support unbiased model training (36,448 transcripts per class). On the other hand, the held-out partition retained the class proportions of the non-redundant set (Table 1).

Beyond primary sequence features used by most classifiers, we computed additional DNA and transcript-associated features that are rarely incorporated in coding-potential tools, focusing on non-B DNA motif and repetitive element features.

Repetitive elements were annotated using RepeatMasker and summarized per transcript after resolving overlaps among transposable elements (TEs), low-complexity regions, and tandem repeats (See Methods and Supplemental Material). For each transcript, we computed 170 repeat-derived features (Table S3), including counts, coverage measures and family-resolved summaries. For these repeat-derived features, we computed both absolute measures (counts/lengths) and length-normalized counterparts (counts per kb and coverage fractions) to mitigate transcript-length confounding. Likewise, non-B DNA motifs were predicted across the GRCh38 genome assembly using G4Discovery [18] and non-B gfa (https://github.com/abcsFrederick/non-B_gfa), yielding 178 motif-derived features per gene body (Table S4). Consistent with previous reports [19], protein-coding and lncRNA sequences in our dataset show distinct distributions of repetitive element content and non-B DNA motifs (Table 2; Table S5), suggesting these features may carry informative signals for transcript classification.

**Table 2.** Descriptive statistics of genomic features for transcripts in the common-CDHIT set.

|  |  |  |  |  |
| --- | --- | --- | --- | --- |
|  | **Protein-coding** |  | **lncRNA** |  |
| | Median ± MAD | | Median ± MAD | |
| **Transcript length** | 1620 |  | 1028 |  |
| | Counts<br>(mean ± std) | % coverage<br>(mean ± std) | Counts<br>(mean ± std) | % coverage<br>(mean ± std) |
| **Rep (Total)** | 1.50 ± 2.55 | 7.12 ± 11.57 | 2.27 ± 2.39 | 28.27 ± 24.35 |
| **NBD (total)** | 288.02 ± 551.34 | 13.63 ± 16.79 | 215.73 ± 415.33* | 13.10 ± 26.96* |
| *For repeats and non-B DNA motifs (NBD), counts indicate the total number of recovered hits (i.e. predicted NBD and number of repeat insertions extracted from RepeatMasker after nested insertion resolution), and coverage represents the percentage of the sequence that is covered by repeats or non-B DNA motifs. Features by repeat class and NBD motif type detailed in Table S5. Acronyms: Rep, Repeats; NBD, Non-B DNA motifs; MAD, Median absolute deviation. \*Since non-B DNA annotation overlaps were not resolved, sequence coverage may be higher than 100%.* |  |  |  |  |

In summary, the “**common-CDHIT”** dataset provides a label-stable (v46–v47 consensus) and redundancy-reduced benchmark, with standardized splits and an extended per-transcript feature compendium, enabling controlled retraining and comparative evaluation of coding-potential classifiers. All transcripts, splits, tool versions/parameters, and derived features are released with a fully reproducible workflow (Methods; Data availability).

### High overall performance but substantial inter-tool disagreement

To assess coding-potential prediction performance on our leakage-controlled benchmark, we surveyed widely used and recently proposed transcript classification tools. We focused on methods with publicly available, functional code that can be retrained from scratch under a controlled protocol, enabling reproducible comparisons on identical training/test splits. We additionally prioritized tools that are actively maintained and collectively represent the major modeling families used for RNA classification.

We selected eight representative tools: lncRNA-BERT [10], CPAT [8], mRNN [20], FEELnc [21], RNAsamba [22], LncFinder [23], LncDC [24], and PLncPRO [25]. They cover transformer-based language models, neural networks, and classical machine-learning approaches (e.g., random forests and gradient-boosted trees). All tools were retrained on the common-CDHIT benchmark using the same cross-validation splits and evaluation metrics (see Methods). We performed 5-fold cross-validation and report out-of-fold predictions, such that each transcript is evaluated exactly once in a held-out test fold, enabling transcript-level agreement analyses across tools.

Across all evaluation metrics, the eight classifiers produced broadly consistent rankings, with lncRNA-BERT, mRNN, and FEELnc forming the top tier. Most tools achieved strong overall performance on this benchmark, with all except CPAT exceeding 80% on balanced accuracy and showing comparably high precision/recall/F1 under the published thresholding rules for each tool (see Methods). Finally, lncRNA-BERT, mRNN and FEELnc consistently reached over 90% across the reported metrics (Fig. 3A; Table S6).

**Figure 3.**
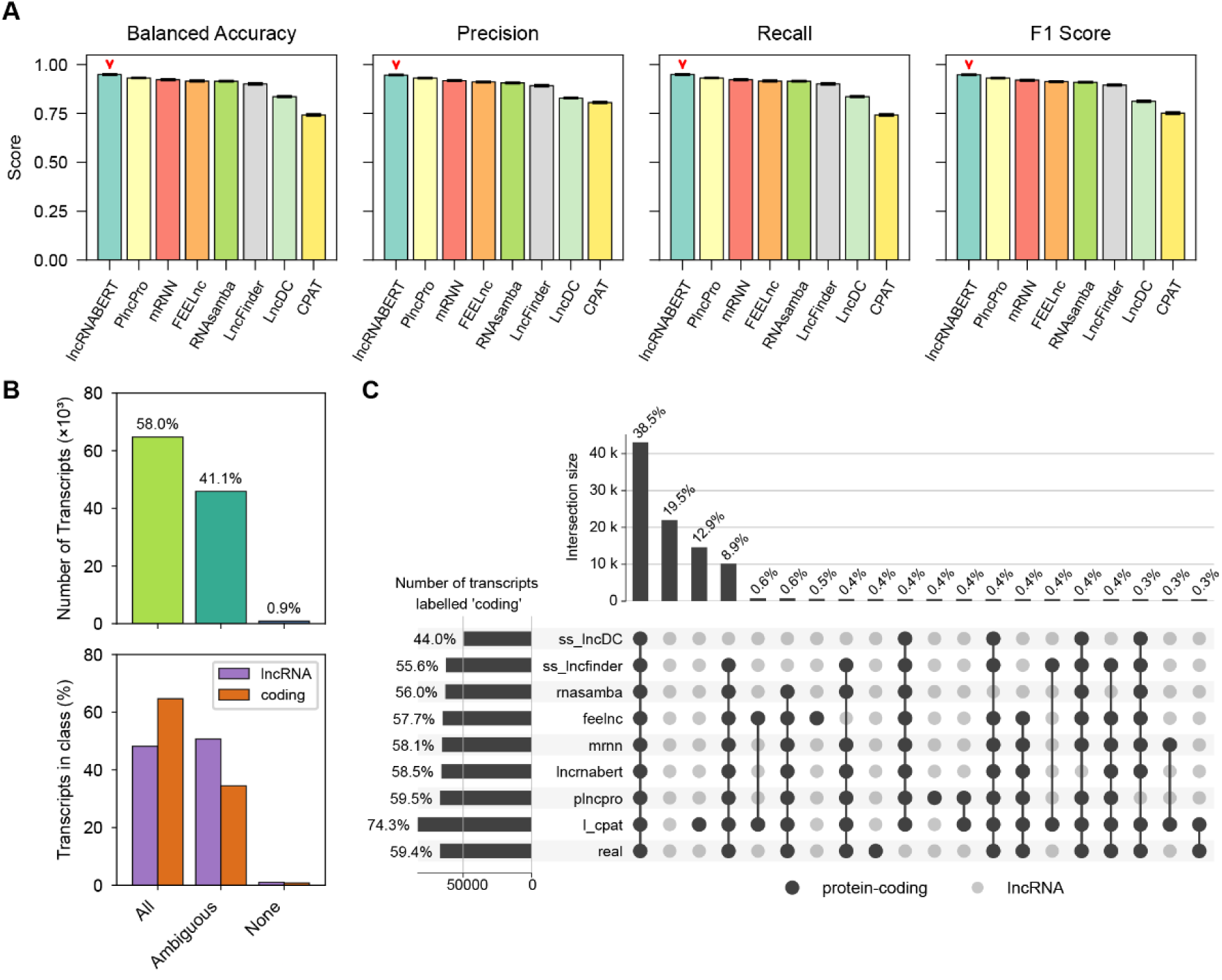
Aggregate performance and inter-tool label agreement. (A) Five-fold cross-validation performance (mean ± s.d.) of the eight retrained models on the common-CDHIT benchmark. Metrics: balanced accuracy, precision, recall, and F1-score (Methods). Average and per-fold values are detailed in Table S6 and Table S7, respectively. (B) Consensus analysis across tools. Transcripts are grouped as unanimously correct (all tools match the reference label), discordant (“Ambiguous”; at least one tool disagrees), or unanimously incorrect (all tools disagree). Left: transcript counts. Right: proportions within each reference class. (C) UpSet plot of top 20 agreement patterns. Black dots indicate the subset of tools predicting “protein-coding” for a given pattern; bars show the number of transcripts assigned “protein-coding” by that exact combination of tools (vertical bars) or by each tool (horizontal bars). Reference labels are from GENCODE.

We next examined inter-tool agreement using the subset of transcripts that produced predictions from all eight tools (N = 111,652; Methods). More than half of transcripts (54.5%) were unanimously correct (all tools matched the GENCODE label), whereas a small fraction (0.9%) were consistently incorrect (all tools disagreed with the label) (Fig. 3B). The remaining 44.6% exhibited discordant predictions, indicating substantial tool-to-tool variability despite high aggregate scores. Stratifying by class, discordance was more prevalent among lncRNAs than protein-coding transcripts (Fig. 3B), consistent with the greater heterogeneity of lncRNA sequences and their more variable annotation status [26].

Agreement patterns further showed that many discordant cases (33.4%) corresponded to near-consensus predictions: transcripts correctly classified by most tools, with one or two tools producing the opposite label (Fig. 3C). In these near-consensus errors, CPAT accounted for a large share of lncRNA misclassifications, whereas PLncPRO and lncDC contributed disproportionately to protein-coding misclassifications (Fig. 3C).

These results highlight a key benchmarking insight: high headline metrics can coexist with substantial disagreement at the transcript level, motivating the analysis of disagreement profiles with transcript resolution. In the next section, we address this by defining transcript “difficulty” in practice, using prediction confidence and inter-tool agreement.

### Transcript stratification using inter-tool agreement and predictive uncertainty

Tools disagreed (produced different class labels) for about 45% of transcripts, with unbalanced results when lncRNA and protein-coding transcripts were considered separately. Moreover, simply counting how many tools disagree on how many transcripts does not provide granular, interpretable, or statistically meaningful insight into these discrepancies. To understand what drives model disagreement and what leads to full consensus on incorrect predictions, we adopted the following strategy: (1) we used an information theory-based decomposition commonly used to summarize ensemble (that is set of 8 selected tools) predictions; (2) we estimated inter-tool disagreement as a disagreement score based on mutual information; and (3) we stratified transcripts into uncertainty groups using top and bottom 10th percentile threshold per coding class (Fig. 4A,B, see Methods). Our final groups contain 11,166 and 3,730 transcripts in the low- and high-entropy groups, respectively (Table 3).

**Figure 4.**
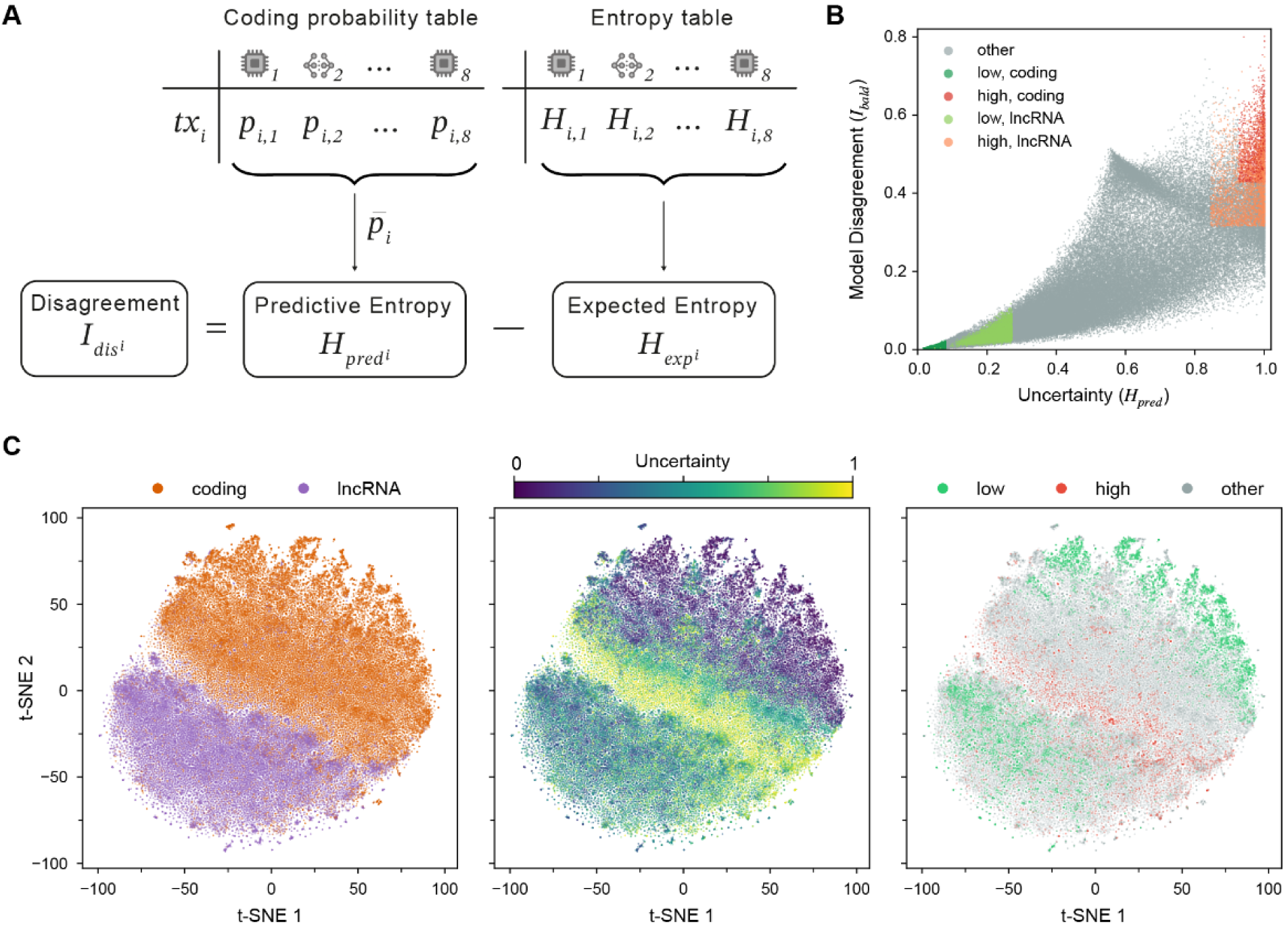
Transcript categorization based on confidence and agreement. (A) Process of entropy calculation. For each transcript, the predicted coding probabilities of eight models are used to calculate the mean predicted coding probability across tools, and the predictive entropy (𝐻_𝑝𝑟𝑒𝑑_) as a measure of classification uncertainty. The expected entropy is calculated as the average entropy of each model’s predicted coding probability, and the disagreement score (𝐼_𝑑𝑖𝑠_) is then derived. (B) Scatter plot of transcripts by ensemble uncertainty and model disagreement. Shades of gGreen represents transcripts with low classification uncertainty (10th percentile, coding: 𝐻_𝑝𝑟𝑒𝑑_ < 0. 080;, lncRNA: 𝐻_𝑝𝑟𝑒𝑑_ < 0. 271 10th percentile) and shades of red highlights transcripts with high classification uncertainty (90th percentile, coding: 𝐻_𝑝𝑟𝑒𝑑_ > 0. 925;, lncRNA: 𝐻_𝑝𝑟𝑒𝑑_ > 0. 844 90th percentile) and model disagreement (90th percentile, coding: 𝐼_𝑑𝑖𝑠_ > 0. 427; lncRNA: 𝐼_𝑑𝑖𝑠_ > 0. 314 , 90th percentile). Transcripts outside these percentile ranges are represented as gray dots. (C) t-SNE visualization of transcripts after dimensionality reduction of the feature table, comprising 124 features computed by the five machine learning models: CPAT (n=4 features) FEELnc (n=8), LncFinder (n=19), LncDC (n=28) and PlncPRO (n=65). Proximity reflects transcript similarity in the original sequence feature space. Plots are colored by coding class (left), predictive entropy (middle) and entropy bin (right).

**Table 3.**
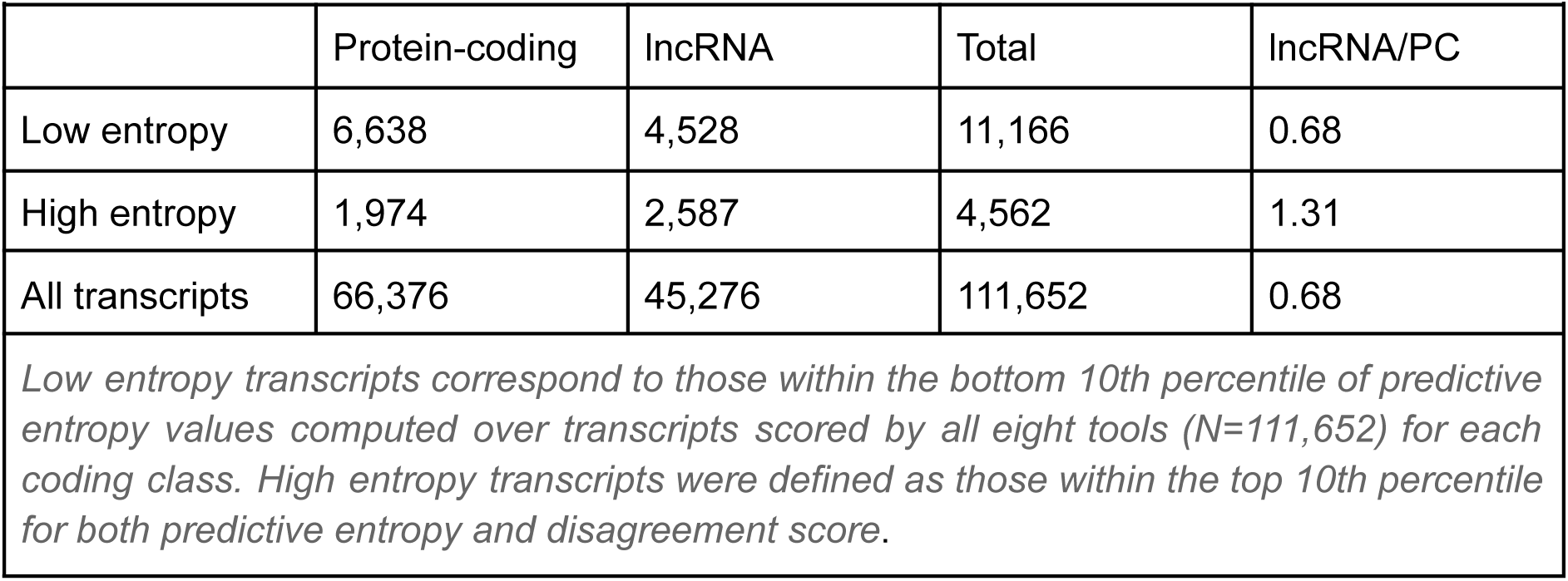
Summary of transcript counts in the low- and high-entropy groups.

To further investigate transcript distribution and contextualize it within their feature space, we performed t-SNE dimensionality reduction using the 124 features extracted by our five ML-based models, and projected transcript labels, entropy values, and uncertainty groups onto this embedding (Fig. 4C). In such visualization, transcript proximity reflects similarity in the original feature space. The class-colored projection revealed a broad but imperfect separation between lncRNA and protein-coding transcripts, with a fuzzy boundary and some transcripts misplaced in the opposing class neighborhood (Fig. 4C, left panel). Entropy values were distributed non-randomly, with low-entropy transcripts clustering tightly within their respective class domains, and high-entropy transcripts localizing at the lncRNA–protein-coding boundary (Fig. 4C, middle panel). This is consistent with the idea that their ambiguous feature profile is a primary driver of tool disagreement. Notably, the low entropy values cluster at the periphery of the protein-coding domain (upper part of the t-SNE) and inside the lncRNA cluster, whereas high-entropy transcripts are mostly located at the frontier between the two classes of transcripts (Fig. 4C, right panel). Closer examination of entropy value distributions revealed that protein-coding transcripts have lower values than lncRNA transcripts (Fig. S2).

The defined low- and high-entropy subsets enable downstream comparisons of feature distributions and misclassification. These subsets provide a benchmark-oriented way to localize where tools agree, disagree, and fail, using a common transcript set.

### Identification of features driving confident classification and discriminating lncRNAs from protein-coding transcripts

Given the observation that high-entropy transcripts localize at the lncRNA–protein-coding classification boundary and that protein-coding transcripts tend to be less entropic than lncRNAs (Fig. S2), we next investigated which features drive this uncertainty, as well as broader lncRNA versus protein-coding discrimination. To identify features driving confident classification and lncRNA-protein-coding differentiation, we focused on the five feature-based (ML) tools, which calculate primary sequence properties (e.g., transcript length, ORF length, k-mer frequencies) and added repeat-based (e.g., class/family diversity, coverage) and non-B DNA motif (e.g., presence of G-quadruplex motifs, etc.) features previously calculated. Deep learning tools were included in agreement analyses but do not expose comparable engineered feature vectors, so they were excluded from feature-attribution analyses.

To assess how strongly each transcript feature associates with distinct entropy and classification groups, we performed univariate statistical analyses paired with effect size estimates (Fig. 5). Since features may share redundant information, we grouped continuous features through hierarchical clustering and retained only the feature with the highest effect size per cluster, with categorical features left as is (see Methods). Transcript length was retained as the representative of its cluster, given its well-documented relevance in coding/non-coding classification. This yielded 155 clusters in total: 66 clusters from ML-based classifier features, 36 from repeat-based features and 53 from non-B DNA features. Complete univariate test results containing non-representative features are shown in Tables S8-S10.

**Figure 5.**
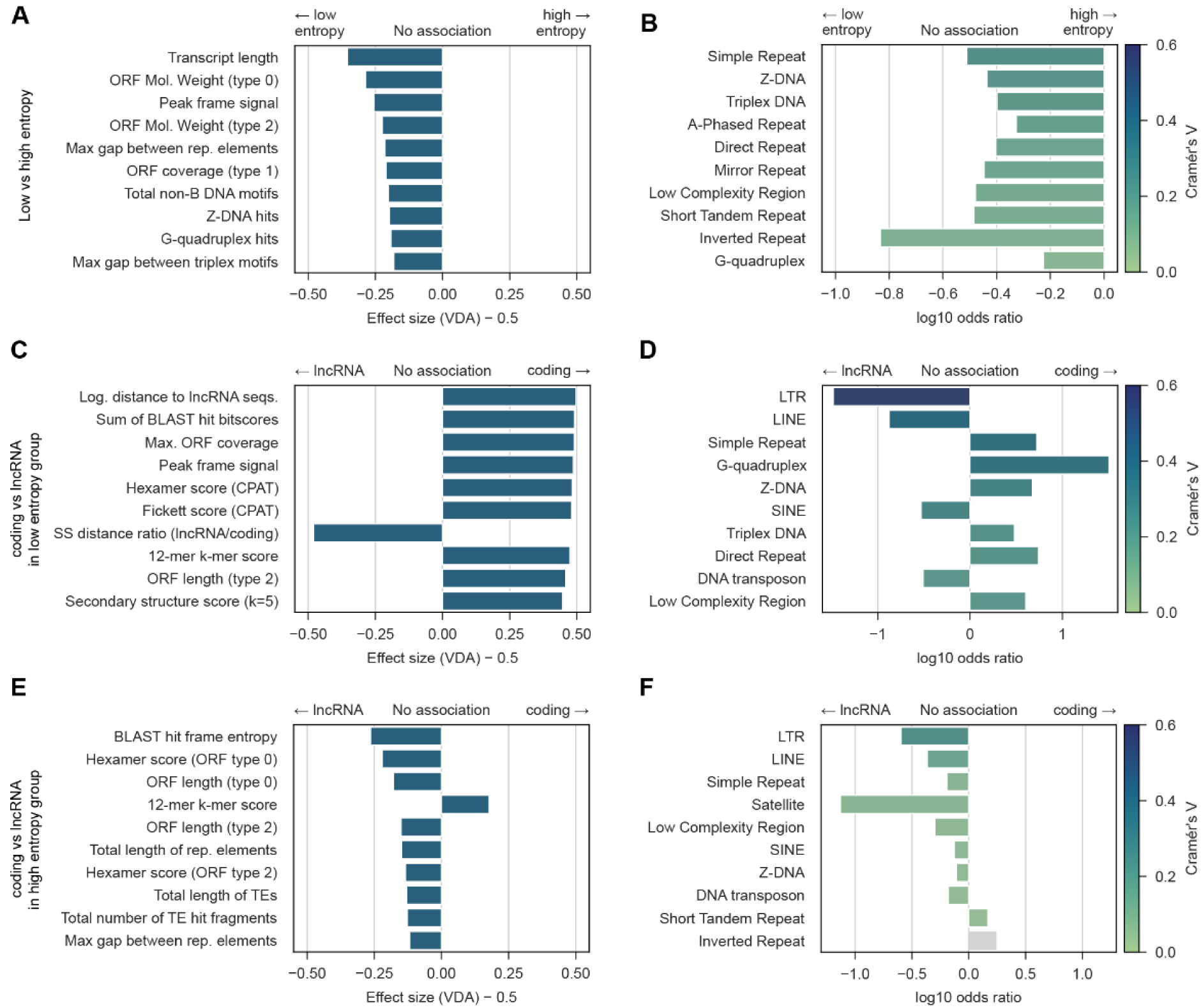
Differential feature associations with entropy groups and transcript class. Feature enrichment testing and prevalence summaries are shown for low- vs high-entropy transcripts (A–B) and for lncRNA vs protein-coding transcripts within the low-entropy (C-D) and high-entropy (E-F) subsets. Statistical tests were selected based on feature type, and all p-values were corrected for multiple testing (FDR; significance threshold p < 0.01). (A,C,E) Top 10 continuous features ranked by effect size (Mann–Whitney U). Effect size of Mann-Whitney U is reported as centered Vargha–Delaney A (cVDA = VDA − 0.5), where 0 indicates no effect and ±0.5 indicates complete separation; the sign indicates association with the left vs right group. Statistically significant features with |cVDA| > 0.1 were considered to have a non-negligible association. Only top feature per feature cluster is reported, except transcript length, which was selected as representative for its cluster. (B, D, F) Top 10 categorical features ranked by effect size (chi-squared). Effect size of chi-squared is reported as Cramér’s V (color scale truncated at 0.6 for visualization; true values in Tables S8-13). Odds ratios indicate direction and magnitude of enrichment. Statistically significant features with V > 0.1 were considered to have a non-negligible association. Non-significant features are colored in gray.

Low- and high-entropy transcripts showed distinct feature associations (Fig. 5A,B). Continuous features such as transcript length, ORF molecular weights and the peak frame signal were significantly higher in low-entropy transcripts (Mann-Whitney U, FDR-corrected p<0.01, centered VDA > 0.15; panel A). Moreover, presence of repetitive elements (simple repeats, low complexity regions) and non-B DNA motifs (Z-DNA, triplex DNA, A-phased, direct, and mirror repeats etc) was also predictive of low-entropy transcripts, with higher motif counts associating with lower entropy. Notably, the maximum gap between either repetitive elements was also associated with low entropy transcripts (Fig. 5A, B).

To further investigate transcripts associated with low and high model uncertainty, we evaluated the differences between lncRNA and protein-coding transcripts within each entropy group (Fig. 5C-F).

In the low-entropy subset, protein-coding transcripts exhibited longer ORFs with higher coverage than lncRNAs (Fig. 5C). Sequence composition metrics, including the Fickett and hexamer scores had larger values in protein-coding transcripts. Interestingly, features derived from non-B DNA motifs and repetitive elements also emerged as discriminative: presence of long terminal repeats (LTR) and long- (LINE) and short- interspersed nuclear elements (SINE) was characteristic of low-entropy lncRNAs, while simple repeats, low-complexity regions and several non-B DNA motifs types were enriched in genes encoding protein-coding transcripts (Fig. 5D). Finally, the secondary structure (SS) distance ratio (a feature summarising how distant SS sequences are to the average) showed that lncRNAs are more dissimilar to the average lncRNA than to protein-coding structures (lncRNA/coding ratio > 1).

On the other hand, feature associations shifted in high-entropy transcripts: lncRNAs displayed longer ORFs and higher hexamer scores, characteristics typically associated with coding potential (Fig. 5E). Moreover, lncRNAs also showed presence of simple repeats and low-complexity regions, previously identified as related to low-entropy coding transcripts (Fig. 5F). This suggests that high-entropy lncRNAs evade confident classification by mimicking coding sequence properties.

Comparing feature associations across entropy groups revealed both consistent and context-dependent patterns. Enrichment of LTR and LINE sequences was associated with lncRNAs in both the low- and high-entropy groups, suggesting that the presence of such elements provides a broadly informative signal for lncRNA classification. In contrast, the 12-mer k-mer score was the only feature with consistently associating higher values with coding transcripts, keeping a robust association with this class across entropy groups.

Overall, statistical enrichment tests showed that: (1) low-entropy transcript features align closely with well-established predictors, validating entropy-based stratification as a reliable post-hoc strategy for curating high-confidence datasets; (2) within the high-entropy group, discriminative signals are substantially weaker and sometimes inverted, reflecting the inherent ambiguity of these sequences; and (3) genomic features underused for transcript classification show concrete and interpretable associations, with LTR presence emerging as a consistent lncRNA signal across entropy regimes. These group-stratified analyses shed light on local drivers of classification uncertainty, but do not assess the global predictive power of individual features. Therefore, we next quantified the importance of these features for classification using multivariate models.

### Identification of global informative features

To identify which features in our new dataset capture global discriminatory signals in a machine learning context, we assessed overall feature importance by examining the learning patterns of Random Forest (RF) models trained on a combined feature dataset. We chose RF because it (1) offers a strong performance–interpretability trade-off for attributing feature contributions and (2) is the same model family used by both PlncPro and FEELnc, the best-performing feature-based tools in our benchmark (Fig. 3; Table S6). Similarly to the univariate analyses, we picked one representative feature per cluster, although this time randomly. We trained one RF model per fold (as defined previously in our cross-validation framework), computed SHAP values for each fold, and averaged them across folds to obtain stable estimates of feature contributions (Fig. 6; Table S14). This approach enabled us to explain global learning patterns by quantifying how each feature influenced transcript classification across the full dataset.

**Figure 6.**
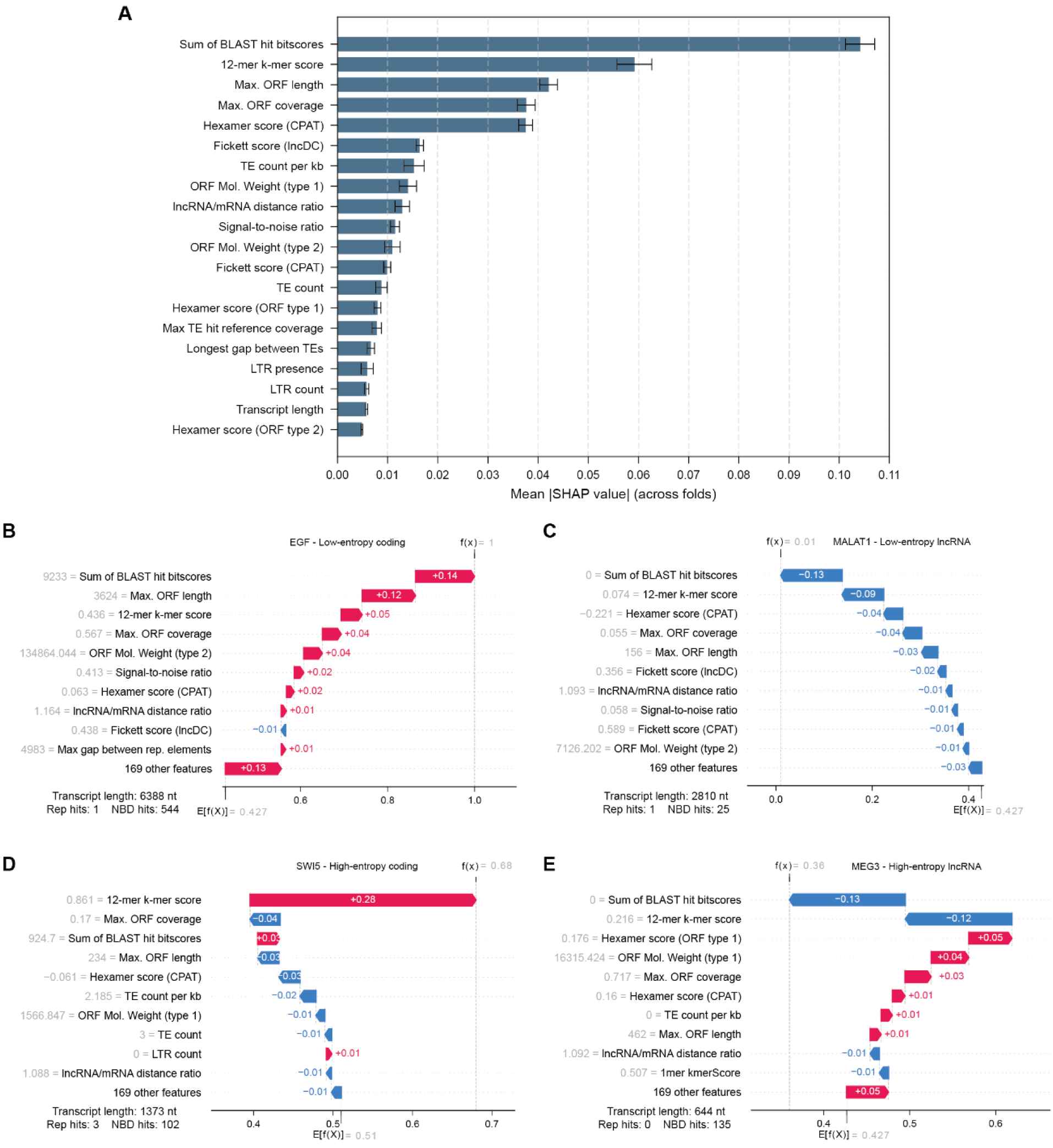
Global feature importance and transcript-level SHAP explanations. (A) Global importance of selected features in the Random Forest model, summarized as mean absolute SHAP value (± s.d.) across cross-validation folds (computed on held-out predictions; see Methods). (B-E). SHAP waterfall plots for representative transcripts showing how individual features shift the model output from the baseline expectation 𝐸[𝑓(𝑥)] to the final prediction 𝑓(𝑥) (coding-class output; reported as probability, Methods). Red bars indicate features increasing the coding prediction (positive SHAP values); blue bars indicate features decreasing the coding prediction (negative SHAP values, toward a lncRNA prediction). Feature values for each transcript are reported on the left of the feature name.

Feature importance analysis based on mean absolute SHAP values indicated that features with stronger contributions to classification were those capturing classical coding-potential information (Fig. 6A). Indeed, sequence composition features (kmer and Fickett scores), together with ORF-related features (coverage, molecular weight) were in the top ten, with four of them showing a stronger impact. The top-ranked feature was the sum of all BLAST hit bitscores, reflecting the cumulative sequence similarity to known proteins as a dominant signal for protein-coding transcripts. Importantly, the second ranked feature was FEELnc’s 12-mer kmerScore, consistent with its behavior as a strong predictor across entropy groups discussed previously.

Interestingly, seven TE-derived features were consistently ranked among informative features, with mean absolute SHAP values between 0.005 and 0.015. While their individual contribution was modest, their collective presence in the top 20 suggests that the repeat content of transcripts carries discriminative information. In this regard, LTR presence stands out, as it was previously identified as a significant predictor in both low- and high-entropy transcript groups. This consistency across low- and high-entropy groups and model-level importance suggests that LTR presence reflects a distinct trait of lncRNAs. Together, these repeat-based features may provide discriminative value beyond classical coding-potential metrics.

To complement the global importance results and provide a feature-based interpretation of misclassification across our entropy-based groups, we examined representative transcripts characterized by distinct confidence and inter-tool agreement. We visualized SHAP values for well-annotated transcripts with available literature and experimental validation, each selected as representative of one group (Fig. 6B-E): (1) a protein-coding transcript from the low-entropy group (EGF), (2) a protein-coding transcript from the high-entropy group (SWI5), (3) a lncRNA from the low-entropy group (MALAT1), and (4) a lncRNA from the high-entropy group (MEG3). We reported the top ten features, and one aggregated term summarizing the contributions of 192 features.

The RF-predicted coding probability of EGF was 1, which classifies it correctly and with high confidence as a protein-coding transcript (Fig. 6B). The primary drivers of “protein-coding” classification of EGF were ORF-related, with a total contribution of +0.20. Remarkably, the longest internal sequence without repetitive elements or TEs contributed an additional 0.04 points to the coding classification (comparable to the 12-mer kmerScore), underscoring the non-negligible contribution of these newly-introduced sequence characteristics. Finally, the residual features accounted for +0.10 of the coding probability, suggesting that multiple features contribute with minimal but additive coding signal. Similar interpretation of the SHAP results for low entropy lncRNA MALAT1 confirmed its correct classification (0.02 coding probability, Fig. 6C), where short predicted ORFs (-0.19) and sequence composition features (Hexamer, K-mer, and Fickett scores, as well as signal-to-noise ratio of EIIP information, -0.13) were the main drivers of non-coding classification. Interestingly, the residual features pushed coding probability of MALAT1 toward 0 with a contribution of equivalent magnitude but opposite sign to that observed in EGF SHAP values, reinforcing the cumulative importance of additional, lower-ranked features.

In contrast, SHAP values of high-entropy, high-disagreement transcripts revealed a consistent pattern. The 12-mer kmerScore contributed towards the correct classification in both examples (+0.18 points for the protein-coding transcript SWI5, and -0.23 points for the MEG3 lncRNA), reiterating the central role of 12-mer composition as clear discriminator of coding-potential. However, a range of other features with opposite contributions substantially reduced classification reliability, yielding near-indifferent predictions: a coding probability of 0.72 for a baseline probability at 0.52 for SWI5 and 0.48 for a decision threshold at 0.548 (Fig. 6D-E).

Beyond the representative cases examined here, the local interpretability of SHAP values of the RF models makes this approach applicable to any transcript of interest, to understand which sequence features drive its classification. Collectively, these analyses revealed the feature profiles underlying inter-tool disagreement. They highlighted discriminative signals exploited by the RF model alongside conflicting features that reduce prediction confidence.

## Discussion

In this study, we aimed to go beyond aggregate performance metrics and characterize where and why tools disagree in protein-coding vs lncRNA classification. By benchmarking eight transcript classifiers with a robust dataset and a controlled protocol, we were able to (1) define transcript groups based on predictive entropy and inter-tool disagreement, (2) investigate feature enrichment across these classes; and (3) assess feature global contributions by training and interpreting RF models. Hypothesizing that limitations of tool’s performances stem, in part, from the difficulty in capturing features beyond standard sequence and ORF-derived signals, we included repetitive elements and non-B DNA motifs, and demonstrated their contribution to transcript classification.

### Model performance and uncertainty

The eight tools we benchmarked showed good overall performance on our dataset, with lncRNA-BERT, mRNN and FEELnc scoring over 90% across balanced accuracy, precision, recall, and F1-score. More than half of transcripts (54.5%) were correctly classified by all tools, but about 45% showed disagreement, especially for lncRNAs where only half were consistently identified correctly compared to 60% for protein-coding transcripts. A small fraction (0.9%) was misclassified by all tools despite high confidence. To dig into this, we used entropy from ensemble predictions to measure uncertainty per transcript, with high entropy means confusion. High-entropy transcripts clustered at the lncRNA/protein-coding boundary in t-SNE embedding, showing they have mixed features that confuse classifiers. Protein-coding transcripts had lower entropy overall than lncRNAs, suggesting lncRNAs are inherently harder to classify due to their diverse sequences and weaker coding signals. This uncertainty global analysis reveals why tools disagree: it’s not just errors, but real biological ambiguity in many transcripts. Finally, transcripts with high confidence but wrong classification suggest they mimic features of the opposite class. Together, these cases justify investigating less common genomic features like repeats and non-B DNA motifs to better capture these differences.

### Feature drivers of uncertainty and (mis)classification

Our entropy-based stratification aimed to clarify why some transcripts are confidently classified while others remain ambiguous. Low-entropy sequences largely conformed to classical protein-coding and lncRNA signatures, with distinctive ORF lengths, k-mer and Fickett scores, as well as characteristic TE and non-B DNA profiles aligning with their reference labels. In contrast, high-entropy transcripts combined discordant features, such as coding-like hexamer composition and ORF metrics in lncRNAs. Thus, these high-entropic transcripts had poor signal dominance, consistent with their position at the lncRNA–protein-coding boundary.

SHAP analyses of the RF models reinforced this view and highlighted where our framework added biological interpretability. As expected, 12-mer, hexamer and ORF-related features remained the strongest global contributors, but TE-derived metrics and non-B DNA attributes, although modest, collectively participated in decision confidence. The four representative transcripts illustrated how both the amount and organization of TE information might modulate classification: EGF and MALAT1 presented opposite extremes of transcript-related size information, supporting clear coding versus lncRNA profiles. In contrast, SWI5 and MEG3 occupied intermediate profiles where substantial TE content coexists with strong coding-potential scores, pulling predictions toward the decision boundary. More broadly, the consistent importance of LTR signals across entropy groups suggests that TEs are not a nuisance for sequence classification but a biologically meaningful axis of lncRNA–protein-coding discrimination, indicating that they might be key contributors to the “ambiguous zone” where current tools disagree.

### Implications for lncRNA biology

Repetitive elements show different patterns between lncRNAs and protein-coding transcripts. Several studies show that lncRNAs associate closely with TEs: Kapusta et al. [27] reported 75% of lncRNAs (GENCODE v13) harbor TE fragments, and Carlevaro-Fita et al. [28] found ∼83% contain TEs. Our RepeatMasker results indicate that 70% of lncRNAs contain at least one repeat, compared to 30% of protein-coding transcripts. In our benchmark, repeat-derived features emerged as informative contributors alongside classical coding signals, supporting a predictive relevance of TE-associated sequence patterns. Beyond simple classification performance, presence of these repetitive elements could be informative of biological function, as suggested by the RIDL hypothesis [19,29]. For example, deleting the Alu-derived SINEB1 domain in MALAT1 disrupts nuclear localization and triggers apoptosis [30]. Our results confirm that LTR and LINE elements are overrepresented in lncRNAs, while simple and non-TE repeats are more prevalent in protein-coding transcripts, suggesting that TE-derived fragments contribute to lncRNA sequence signatures that can help separate them from coding transcripts. Interestingly, TEs inserted in lncRNA exons showed increased divergence from the consensus sequence. Beyond carrying predictive information, this is consistent with the reduced evolutionary constraints of non-coding transcripts [19].

Non-B DNA motifs leading to unusual secondary structures encode regulatory information in genes beyond primary sequence. Non-B DNA structures in the body of genes, including genes transcribing lncRNAs, affect replication [16,31], and transcription initiation and progression. RNA G-quadruplexes in lncRNAs also suggest regulatory roles [32]. In our analyses, non-B DNA motif features (G-quadruplexes, Z-DNA, and triplex motifs) were overrepresented in the gene bodies of low-entropy transcripts, which are derived from protein-coding genes. This asymmetry is consistent with a model in which protein-coding gene bodies are enriched for non-B DNA motifs (based on observed low entropy, Figure 5D), whereas lncRNAs more often carry RNA motifs (e.g., recognition and binding domains) that recognize and bind these structures. It also suggests that while non-B DNA structures are retained at protein-coding loci under evolutionary constraint, lncRNAs rely more on RNA-level architecture (secondary/tertiary structure and multivalent protein binding) to achieve regulatory specificity. Indeed, lncRNAs mainly function through secondary/tertiary structure, which specifies which proteins, RNA or DNA regions lncRNA can bind.

High-entropy lncRNAs may reflect younger evolutionary age or lower conservation. One possible explanation for the high-entropy profile is reduced evolutionary constraint, leading to weaker or more conflicting sequence signals and increased inter-tool disagreement. Entropy could proxy lncRNA age with higher entropy for younger or less conserved lncRNAs, more disordered sequences under relaxed selection. This hypothesis can be tested using PhyloCSF [6] or phastCons [33] scores on high- vs. low-entropy groups in future studies.

### Limitations and future directions

The model ensemble included eight classifiers; however, feature importance analyses were restricted to the feature-based models due to the inherent interpretability challenges of DL models. Further work integrating attention weights from the DL models would help bridge this gap. Accordingly, our modular pipeline architecture allows the inclusion of additional tools, extending the ensemble with future methods as the state of the art evolves.

A related concern is the stability of the entropy group assignments upon changes in ensemble composition. Defining these groups by using thresholds on entropy values instead of vote-count agreement limits the impact of ensemble variation. Future analyses could treat entropy as a continuous response variable in a regression framework, which would allow for a more granular characterization of features predictive of classification uncertainty.

Our framework relies on quantifying prediction disagreement across ensemble members based on sequence-derived features, pinpointing transcripts for which the sequence information alone is insufficient to confidently assign a class. Importantly, this could be useful to flag transcripts for experimental validation or reannotation. However, model uncertainty should not be interpreted as biological ambiguity. Coding potential is inherently context-dependent: lncRNAs frequently show tissue-specific expression, some undergo condition-specific translation through tightly regulated ribosome recruitment, and dual-function transcripts with both coding and non-coding roles blur the boundary between classes. As a result, such transcripts may be biologically ambiguous yet confidently classified by our ensemble, or vice versa.

While the current set of features has proven useful for high-performing models, it exhibits high multicollinearity, with many features correlated with transcript length. This can affect interpretability: when globally important features are highly correlated, feature-importance rankings may distribute importance across redundant predictors and can obscure characteristics that matter for specific outlier transcripts or borderline cases. Moreover, SHAP-based interpretation depends on the choice of explanatory model (here a Random Forest), which can itself be influenced by collinearity in the feature space. Exploring alternative interpretability frameworks and model classes less sensitive to correlated predictors may help reveal complementary signals.

Finally, our analyses incorporated repetitive elements evaluated exclusively at the transcript-level. While this is a natural scope when evaluating classifiers that operate on mature transcript sequences, TEs are also abundantly distributed across introns. These intronic TEs may also influence transcript biology from transcriptional regulation to RNA maturation [34,35]. Integrating such intronic features, together with additional layers of genomic context, might provide further insights. Furthermore, while our analysis of non-B DNA motifs was conducted at the genomic level, structural predictions on RNA sequences could be explored to capture potential non-canonical RNA conformations that might influence RNA stability, localization, or translation. Indeed, recent work has demonstrated the value of alternative genomic-context features in coding/non-coding classification [36]. Future work could include positional information (intergenic, antisense or overlapping regions), exon-intron structure, chromatin accessibility, and phylogenetic conservation metrics to better capture evolutionary and regulatory constraints on transcripts structure, regulation and function.

## Conclusions

Our benchmarking strategy helps elucidate transcript profiles that drive misclassification and inter-tool disagreement in protein-coding versus lncRNA prediction. It serves two main purposes: (1) highlight transcript properties associated with coding versus non-coding types, and (2) identify feature families that can improve the robustness and interpretability of future classifiers.

Our study further shows that benchmarking can be informative beyond aggregate performance metrics, by revealing transcript-level agreement patterns and systematic characteristics of “easy” versus “difficult” cases. Refining this framework will support more interpretable and biologically informed models, and provide a clearer view of the heterogeneous sequence profiles that underlie lncRNA annotation.

A practical implication for genome annotation workflows is to use inter-tool agreement/entropy as a quality-control step: transcripts with high agreement and low entropy can be considered high-confidence predictions, whereas high-entropy or high-disagreement transcripts should be explicitly flagged as uncertain. These flagged cases can then be prioritized for orthogonal evidence (e.g., ribosome profiling/proteomics support for coding potential or evolutionary conservation), rather than being treated as definitive labels. Indeed, flagged transcripts with ambiguous features may actually hide dual functions, both at the RNA level and as carriers of micro-ORFs with potential to encode small peptides. The complicated ambiguous classification of high-entropy transcripts suggests that the distinction between coding and non-coding sequences is rather a spectrum than a strict boundary. Altogether, our results motivate uncertainty-aware, feature-rich classifiers and a community practice of propagating prediction confidence into downstream annotation and validation. Beyond improving computational workflows, our study contributes to refining the biological definition of lncRNAs.

## Methods

This study consists of four main components: (1) constructing a label-stable, redundancy-reduced benchmark dataset with extended genomic features (common-CDHIT); (2) benchmarking eight lncRNA/mRNA classifiers under a standardised 5-fold cross-validation framework, producing out-of-fold coding probabilities and binary labels; (3) characterising transcript-level classification uncertainty and inter-tool disagreement using entropy-based metrics, and stratifying transcripts into distinct uncertainty regimes; and (4) performing feature importance analyses across complementary feature sets to identify the sequence features driving classification behaviour. Each component is described in the sections below. Additionally, the Fig. S1 summarizes the general workflow of the study, including dataset pre-processing, model training and evaluation as well as downstream analyses.

### Construction of the benchmarking dataset

Dataset construction involved three steps: (1) selecting transcripts with stable protein-coding or lncRNA biotype assignments across GENCODE releases v46 and v47; (2) applying sequence-level redundancy reduction using CDHIT to prevent data leakage; and (3) computing an extended set of genomic features (repetitive elements and non-B DNA structural motifs) to complement sequence-intrinsic features of the benchmarked tools. The resulting dataset is referred to as the common-CDHIT set.

#### 1. Label-stable GENCODE dataset

We downloaded human transcript sequences and annotations from the GENCODE project (releases v46 and v47) based on the GRCh38 genome assembly. Transcripts were classified into two main categories: protein-coding transcripts and lncRNA, based on the GENCODE FASTA files (e.g., gencode.v46.pc_transcripts.fa and gencode.v46.lncRNA_transcrips.fa) and the respective files for each release. Transcripts outside these categories were discarded for our analyses.

We then generated a dataset containing only transcripts present in both releases v46 and v47 with no change in class label. We refer to this subset as the “common dataset”. This cross-release filtering focuses model training and evaluation on transcripts with stable annotations, improving the reliability of downstream analyses.

#### 2. Redundancy reduction of transcripts

To minimize sequence similarity within and between training and testing sets and prevent overfitting, we surveyed the strategies used by the original publications of the tools used in the study and followed the most used approach. Clustering was performed on the “common dataset” using CD-HIT with default parameters (90% sequence identity threshold, 90% coverage, word size of 8 nucleotides) [17]. For each cluster, we retained only the representative sequence, creating a non-redundant dataset “common-CDHIT” for downstream analyses.

#### 3. Computation of genomic features

##### Repetitive element analyses

Transcripts in the common-CDHIT dataset were scanned for repetitive elements using RepeatMasker v4.2.1 [37] in sensitive mode with the default human Dfam library (v3.9). When overlaps occurred, overlapping annotations were resolved by retaining high-confidence hits based on alignment scores and annotation quality (see Supplementary Methods). Retained hits were separated in three major classes: (1) transposable elements, (2) pseudogenes, and (3) simple repeats and low complexity regions. Then, each class was processed using a custom hierarchical feature extraction pipeline (https://github.com/cbib/rep_extraction_pipeline), which computed 170 manually curated features at three levels: (1) individual hits (length, divergence, indels, alignment scores), (2) aggregated elements accounting for hit fragmentation (total length, fragment count, class and family assignment), and (3) transcript-level summaries (total hit counts, total coverage, hit counts per family, presence/absence flags, and inter-hit statistics). Finally, feature dimensionality was reduced through a hierarchical clustering approach described above.

##### Non-B DNA prediction

Non-B DNA motifs were predicted on the GRCh38 reference assembly using the *non-B_gfa* (gfa) pipeline (https://github.com/abcsFrederick/non-B_gfa), generating genomic intervals for A-phased repeats (APR), direct repeats (DR), inverted repeats (IR), mirror repeats (MR), short tandem repeats (STR), and Z-DNA. Triplex-forming motifs (TRI) were defined as the subset of MR predictions flagged as triplex-prone by the gfa output. G-quadruplexes (G4) were predicted on GRCh38 using G4Discovery [18] with default parameters, reporting both pqsfinder [38] and G4Hunter [39] scores. All non-B DNA motif calls were intersected with GENCODE v47 gene intervals using BEDtools to assign motif occurrences to genes encoding transcripts in the common-CDHIT dataset (Supplemental Information).

For each transcript, we computed 178 non-B DNA features, including per-motif-type presence/absence, hit counts, length and gap statistics, as well as global summary features (e.g., diversity, coverage) calculated on the full gene bodies(exons+introns). Because a locus can support multiple non-B DNA structural annotations, overlapping motif calls were retained (no overlap resolution) when computing features. Feature extraction was performed using a custom pipeline (https://github.com/cbib/nbd_extraction_pipeline).

### Benchmarking protocol

The benchmarking workflow consisted of: (1) selecting retrainable tools with public, runnable code; (2) retraining each tool under the same 5-fold cross-validation splits; (3) producing coding scores and binary labels for out-of-fold transcripts; and (4) evaluating global classification performance with macro-averaged metrics. These steps are detailed below.

#### 1. Selection of retrainable tools

Selection of tools for the benchmark was guided by five criteria:

● public source code availability,
● compatibility with custom retraining,
● capacity to yield soft predictions,
● algorithmic diversity,
● support for feature-based interpretability.

These criteria are further detailed in Supplemental Methods. The final selection of tools is detailed below in Table 4.

**Table 4:**
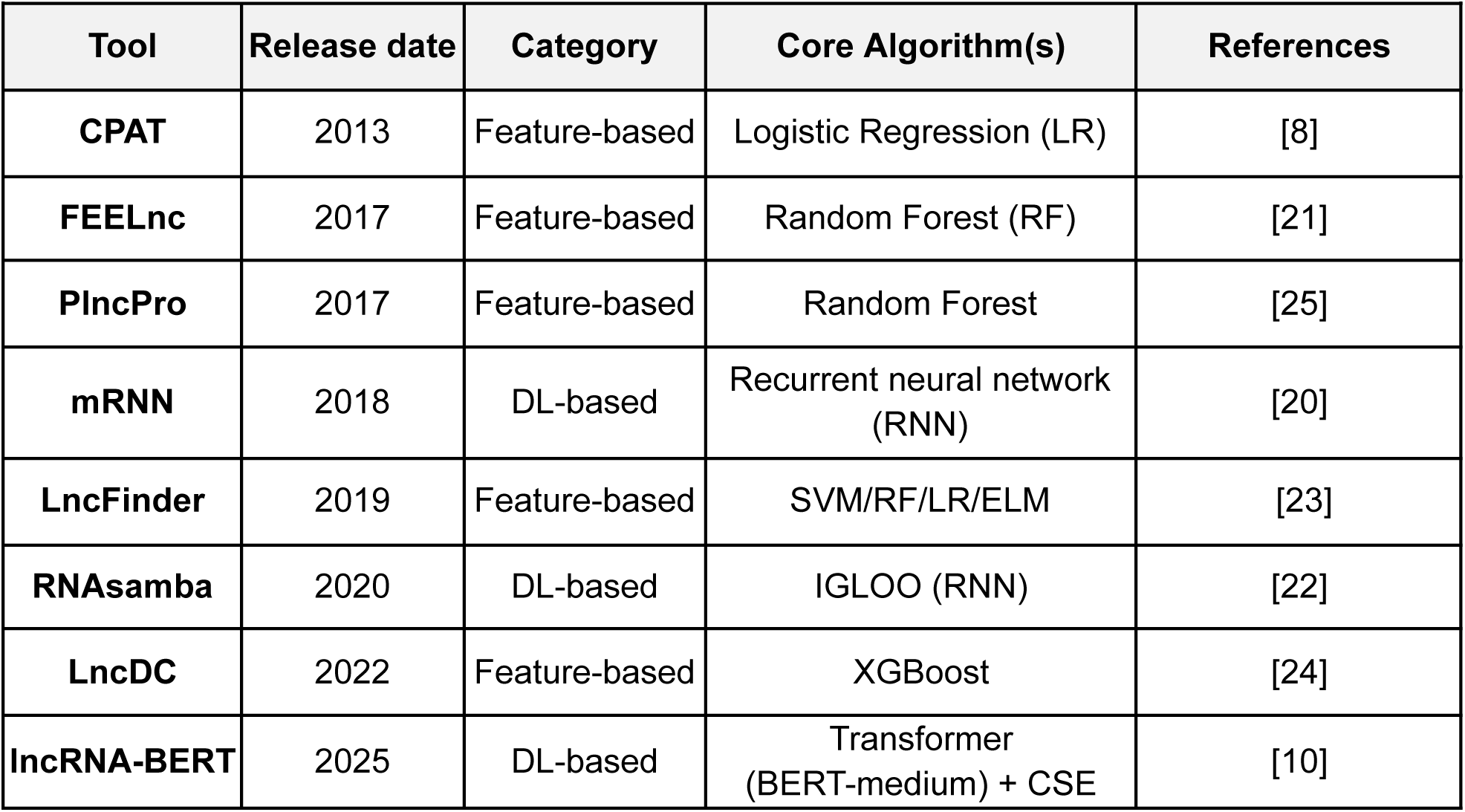
Tools selected for benchmarking.

Technical details about the selected models are available in Supplemental Methods. For tools that proposed several training methods, we selected the one displaying better classification performance. Finally, we ensured tool predictions were diverse enough to be able to analyze classification differences (Figure S1). All tools were used for comparison analyses (see uncertainty and agreement analyses). However, note that feature analyses were only performed using the five feature-based tools (i.e., CPAT, FEELnc, PlncPro, LncFinder and LncDC).

#### 2. Cross-validation setup and model training

All tools were trained following the same benchmarking setup. The common-CDHIT dataset was split in 5 cross-validation folds, where 20% transcripts from each class were held-out as a fixed test set per fold. Given the class imbalance in the original dataset, all transcripts belonging to the minority class (i.e. all lncRNAs) were retained in the training set. To avoid training bias due to the higher prevalence of protein-coding sequences, a random sample of equal size was drawn from the remaining protein-coding transcripts. Additionally, for tools incorporating a validation step during model training, a subset of training transcripts was held-out for this purpose, following each tool’s published recommendations (Supp. Methods) The eight models were trained using the original protocols, using either (1) scripts based on the tool libraries (e.g., LncFinder), (2) original source code with minimal modifications (e.g., LncDC, mRNN) or (3) directly executable training scripts provided by the authors. Exact parameters for each tool are detailed in Supplemental Methods.

#### 3. Coding scores and label assignment

Transcript classification was performed independently for each tool and fold. Each model produced both a coding probability score (ranging from 0 to 1) and a binary classification label (coding / non-coding). Decision thresholds were established according to published procedures, with most tools applying a default 0.5 probability cutoff. For tools using custom thresholds, we recalculated them accordingly per fold before determining the classification label. For tools that define lncRNAs as the positive class, coding scores were derived by inverting the non-coding scores. A summary of model probabilities and entropies, reflecting how confident model predictions are, is provided in Fig. S2.

##### Classification results aggregation

Due to constraints imposed by certain benchmarked tools (e.g., minimum or maximum transcript length, absence of identified ORF, presence of non-canonical nucleotides), 612 transcripts could not be assigned a label by all classifiers. These transcripts were excluded from downstream analyses requiring classification scores from all tools, resulting in a final dataset of 111,652 sequences with associated features, coding probabilities and classification labels. This set of classified transcripts was used for both global performance evaluation of models, entropy analyses and feature importance.

#### 4. Classification performance evaluation

Global model performance was evaluated using standard metrics: balanced accuracy, precision, recall and F1-score. Given the class imbalance, macro-averaged scores were computed. Detailed formulas for the scores are further detailed in Supplemental Methods.

### Uncertainty and agreement analyses

Our 8-tool ensemble was used to investigate transcript-level correctness, agreement, and uncertainty using entropy-based summary statistics of the predicted coding probabilities.

Specifically, we (1) computed predictive entropy, which quantifies uncertainty in the ensemble mean prediction; (2) produced an inter-tool disagreement score, which captures variability across tools; and (3) stratified transcripts into different uncertainty groups for downstream comparisons.

#### 1. Uncertainty

Transcript classification uncertainty was quantified using Shannon entropy of the mean predicted coding probability across tools. For each transcript, coding probabilities were averaged across tools to compute the mean posterior probability for the coding class (𝑝_𝑐𝑜𝑑𝑖𝑛𝑔_). Shannon entropy was calculated as:

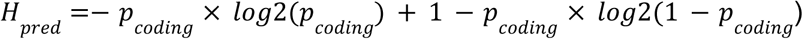

This metric produces values from 0 to 1, where 𝐻 = 0 corresponds to extreme mean probabilities (near 0 or 1) indicating complete certainty (all tools agree with high confidence) and 𝐻 = 1 indicates maximum uncertainty of the ensemble mean 𝑝_𝑐𝑜𝑑𝑖𝑛𝑔_ ≈ 0. 5 (average model prediction is uncertain). Classification entropy distributions per model and ensemble predictive entropy are shown in Figure S3B-C.

#### 2. Disagreement

Inter-tool disagreement was summarized using an entropy-gap score. We first computed the expected entropy of each transcript as the average entropy of individual model predictions:

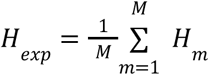

where 𝐻_𝑚_ represents Shannon entropy of the coding probability of a transcript given by a model 𝑚. Model disagreement was then defined as the difference between predictive and expected entropy.

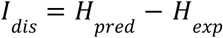

Low 𝐼_𝑑𝑖𝑠_ values indicate high ensemble consensus, while high 𝐼_𝑑𝑖𝑠_ values indicate substantial model disagreement.

#### 3. Uncertainty-based transcript stratification

We categorized transcripts into uncertainty groups using a percentile-based approach. We split all predicted transcripts (111,652) in coding and non-coding and computed for each group the 10th and 90th percentiles of our uncertainty metric (𝐻_𝑝𝑟𝑒𝑑_) and the 90th percentile of the disagreement metric (𝐼_𝑑𝑖𝑠_ ). Transcripts with 𝐻_𝑝𝑟𝑒𝑑_ < 10th percentile were assigned to the “low entropy” group, while sequences with 𝐻_𝑝𝑟𝑒𝑑_ and 𝐼_𝑑𝑖𝑠_ > their respective 90^th^percentiles were assigned to the “high entropy” group. This grouping strategy enabled us to identify transcripts with distinct prediction difficulty profiles and assess systematic differences in biological features across uncertainty groups. These differences were evaluated using univariate statistical tests described below.

### Feature-based analyses

From the eight models benchmarked here, five are machine learning tools based on engineered transcript features. We generated the corresponding feature tables and assembled a unified feature matrix (ML set). Additionally, we included repeat-derived features (Rep set) and Non-B DNA motif features (NBD set) features computed as described above (see Supplemental Material for additional details).

#### 1. Feature processing

We applied the following preprocessing steps (1) removal of constant features (i.e., those with the same value across all transcripts); (2) separation of categorical and continuous variables; (3) estimation of continuous feature redundancy by calculation of a Spearman correlation matrix; (4) clustering of correlated features; (5) selection of a random representative feature per cluster; and (6) combination of representative continuous features with categorical features encoded as binary indicator variables via one-hot encoding). The number of features after each processing step is detailed in Table S15.

### Clustering and representative feature selection

Specifically, continuous features were grouped using hierarchical clustering (Ward linkage) on a Spearman correlation-derived distance matrix. The number of clusters was chosen by maximizing the silhouette score over a grid of cut thresholds (Fig. S2), and one feature was randomly selected per cluster as representative

### Dimensionality reduction for feature space exploration

To visualize the feature space of transcripts classified by the benchmarked models, t-SNE embeddings were computed using the scikit-learn Python library with 1000 iterations and a perplexity of 30. The input feature space comprised all 128 numerical, non-constant engineered features from the 5 feature-based models benchmarked in this study: CPAT (N=4 features) FEELnc (N=8), LncFinder (N=19), LncDC (N=28) and PlncPRO (N=69).

### Univariate statistical testing of transcript features

Differences in continuous features between transcript groups (coding class or entropy groups) were assessed using Mann-Whitney U test, with effect size quantification by the Vargha-Delaney A (VDA) statistic. For visualization, we report a centered effect size (VDA − 0.5), where 0 indicates no effect and ± 0.5 indicates complete separation between groups. Differences in categorical feature prevalence were assessed using the chi-squared test of independence, with effect size quantified by Cramér’s V statistic and the directional magnitude expressed in odds ratios (OR). To account for multiple comparisons, p-values were corrected using the Benjamini-Hochberg method. Results with adjusted p-value lower than 0.01 were considered statistically significant.

#### 2. Global feature importance

A Random Forest model was then trained for each of the 5-fold train sets on the selected features, comprising 155 representative continuous features and 24 categorical features. SHAP values were then computed for the held-out set of each fold to obtain transcript-level feature explanations (waterfall plots). Importantly, SHAP values are expressed directly in probability space, such that for each transcript, the sum of all feature SHAP values and the baseline (mean predicted probability assigned by the RF classifier) equals the coding probability assigned by the RF classifier. Finally, SHAP values from all held-out sets were combined to obtain global feature importance summaries (mean absolute SHAP value across all transcripts). The classification performance metrics of the trained RF classifiers are detailed in Table S16.

### Development and computation tools

Dataset creation, model training and data analyses were implemented as a Snakemake pipeline (v9.13.2) [40], using Python and R. Separate conda environments were created for each model to ensure reproducibility and portability. A dedicated analysis environment was used for data processing and visualisation, including Biopython and gffutils (biological data handling); numpy and pandas (data processing); scikit-learn, scipy and shap (statistics and dimensionality reduction), and matplotlib/seaborn/plotly/upsetplot (visualization). Environment configuration is available through environment files in the GitHub repository (https://github.com/cbib/uncertainty_aware_lnclassifier).

All computations were performed on the University of Bordeaux high-performance computing (HPC) infrastructure operated by the Center of Bioinformatics of Bordeaux (CBiB). GPU-accelerated tasks (RNAsamba and lncRNA-BERT training/inference) were run on NVIDIA H100 GPUs.

## Abbreviations

## Declarations

### Ethics approval and consent to participate

“Not applicable”

### Consent for publication

“Not applicable”

### Availability of data and materials

Data produced during the analyses is deposited in Zenodo https://doi.org/10.5281/zenodo.19551649

## Authors’ contributions

DGR, DC and MN designed the benchmarking protocol. DGR, MG, RB and SKM prepared the benchmarking datasets. DGR implemented the reproducible Snakemake pipeline and performed model retraining and evaluation. DGR and MG analyzed the data. DGR, MG, MN and DC interpreted the results. DGR drafted the manuscript; DGR, SKM, KDM, MN and DC reviewed and edited the manuscript. MN, DC and KDM supervised the study and acquired funding. All authors read and approved the final manuscript.

## Acknowledgements

We gratefully acknowledge support from the CBiB (Centre de Bioinformatique de Bordeaux) of the University of Bordeaux for the computing resources. We are also thankful to all the members of the CB&B team for their scientific discussions.

