## Supplemental Information for "Uncertainty-aware benchmarking reveals ambiguous transcripts in mRNA–lncRNA classification"

### Benchmarking dataset

#### *Supplemental information*

We performed a systematic comparison between consecutive GENCODE versions (v46 and v47 for human). For each transcript, we tracked its presence and classification status across versions, categorizing transcripts into groups: (1) common transcripts with stable classification (here called “common dataset”), (2) common transcripts with changed classification, (3) newly added transcripts with assigned classification and (4) newly added transcripts without assigned classification. Additional groups were: (5) common transcripts that lost classification, (6) common transcripts that gained classification, (7) common transcripts without classification, (8) removed transcripts with classification, (9) removed transcripts without classification. Groups 2 to 9 were excluded from the current analyses.

#### *Supplemental methods*

##### Coding sequence extraction

The benchmarked classification tools CPAT, lncDC and LncFinder required coding sequences (CDS) of transcripts as training inputs. For protein-coding transcripts in the training set, we extracted their corresponding using the Ensembl CDS annotations.

#### *Supplemental tables*

**Table S1:** Transcript classification tools displayed in Figure 2A.

| Tool | Year | Reference |
|---|---|---|
| CPC | 2007 | Kong et al., 2007 [1] |
| PORTRAIT | 2009 | Arrial et al., 2009 [2] |
| PhyloCSF | 2011 | Lin et al., 2011 [3] |
| CPAT | 2013 | Wang et al., 2013 [4] |
| CNCI | 2013 | Sun et al., 2013 [5] |
| lncRScan | 2015 | Sun et al., 2015 [6] |
| COME | 2017 | Hu et al., 2017 [7] |
| FEELnc | 2017 | Wucher et al., 2017 [8] |
| CPC2 | 2017 | Kang et al., 2017 [9] |
| PLncPRO | 2017 | Singh et al., 2017 [10] |
| longdist-SVM | 2017 | Schneider et al 2017 [11] |
| mRNN | 2018 | Hill et al., 2018 [12] |
| lncRNAnet | 2018 | Baek et al., 2018 [13] |
| LncADeep | 2018 | Yang et al., 2018 [14] |
| CPPred | 2019 | Tong et al., 2019 [15] |
| PredLnc | 2019 | Liu et al., 2019 [16] |
| LncFinder | 2019 | Han et al., 2019 [17] |
| RNAsamba | 2020 | Camargo et al., 2020[18] |
| LncDC | 2022 | Li et al., 2022 [19] |
| LncRNA-BERT | 2025 | Romejin et al., 2025 [20] |

| **Table S2:** Number of transcripts by class and version, as displayed in Figure 2B. | | |
|---|---|---|
| **From (v46)** | **To (v47)** | **Count** |
| new transcripts | lncRNA | 131,424 |
| protein-coding | protein-coding | 111,838 |
| no class | no class | 82,231 |
| lncRNA | lncRNA | 59,674 |
| new transcripts | protein-coding | 331 |
| lncRNA | removed transcripts | 187 |
| new transcripts | no class | 52 |
| lncRNA | no class | 43 |
| no class | protein-coding | 26 |
| lncRNA | protein-coding | 23 |
| protein-coding | removed transcripts | 20 |
| no class | removed transcripts | 11 |
| protein-coding | no class | 9 |
| no class | lncRNA | 7 |
| protein-coding | lncRNA | 1 |
| | **Total** | **385,877** |

### Classification tools: selection, retraining and performance

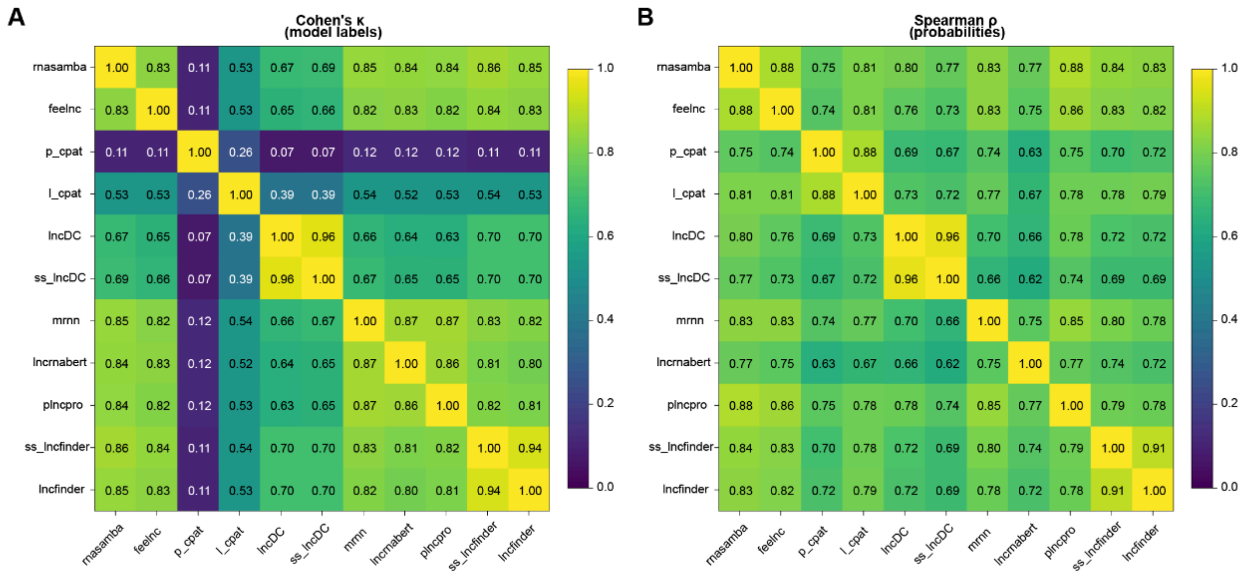

#### *Supplemental figure*

**Figure S1. Diversity of model predictions.** (A) Heatmap representing Cohen's kappa statistic on the similarity of predicted labels. (B) Heatmap representing spearman correlation coefficient of model coding scores.

#### *Supplemental methods*

##### Tool's specific methods

First, tools were required to have publicly available source code, with a preference for widely used and actively maintained methods, ensuring the reproducibility and long-term accessibility of the benchmark. Second, inclusion required support retraining directly or with minimal modifications, allowing all models to be evaluated with any custom benchmarking dataset. Third, since uncertainty analyses are grounded on approximating model scores to coding probabilities, benchmarked tools must report classification as a continuous score between 0 and 1. Fourth, selected models span a diversity of algorithm families (including logistic regression, support vector machines, tree algorithms, neural networks and transformer-based architectures) so that model disagreement reflects real uncertainty rather than simple correlated errors arising from methodological similarities. Finally, a substantial number of these architectures were required to be feature-based, providing the basis for post-hoc interpretability analyses of the discriminative features in different uncertainty regimes.

##### *CPAT*

The Coding Potential Assessment Tool (CPAT) is a logistic regression model that uses four sequence-based features to classify transcripts [4]. These features are (1) maximum ORF length, (2) ORF coverage (or ORF-to-transcript length ratio), (3) Fickett Score (a metric summarizing codon usage distribution) (Fickett, 1982), and (4) hexamer score (usage bias of adjacent codons). The original model was trained with 10 000 protein-coding transcripts from the RefSeq database [21] and 10 000 lncRNAs from Gencode [22]. The CPAT model was evaluated by 10-fold cross-validation, and a probability cutoff of 0.364 was selected to maximize both sensitivity and specificity when labeling transcripts. Finally, classification performance was tested on an independent set containing 4000 protein-coding transcripts from RefSeq and 4000 lncRNAs from a different curated dataset (Cabili et al 2011), achieving a sensitivity of 0.96 and a specificity of 0.97.

We used CPAT version 3.0.5 to train a custom model following the instructions in the original publication and the online documentation. Since version 3.0.0, CPAT allows users to select the ORF with highest coding probability instead of the longest ORF for ORF feature calculation. We therefore trained models with both options and selected the best-performing one (Table S6). A reference hexamer frequency table was built using only the training dataset as inputs. We ran a 10-fold cross validation and found an optimal probability cutoff value of 0.410, slightly higher than the original 0.364. The final logistic regression model was trained on the full training dataset and model performance was evaluated on the common test set after assigning classification labels based on the cutoff (i.e., p < 0.410 -> lncRNA; p >= 0.410 -> protein-coding).

##### *FEELnc*

The Flexible Extraction of LncRNAs (FEELnc) tool is a pipeline for lncRNA detection and annotation [8]. It includes the FEELnc codpot module (hereon called simply FEELnc) which is a Random Forests (RF) model that calculates a coding potential score based on three sequence features: (1) transcript length, (2) ORF coverage and (3) Kmer Scores (summarizing multi kmer frequency differences between coding and non-coding transcripts). The transcript set used for training FEELnc was extracted from Gencode v24 and was composed of 5000 coding transcripts (those with biotype 'protein_coding') and 5000 lncRNA transcripts (with biotypes 'lincRNA' and 'antisense'). A held-out test set was also created using 5000 coding and 5000 lncRNA transcripts. Importantly, only one transcript per gene was used, thereby avoiding sequence redundancies within and between training and testing datasets. The method for extracting ORFs and the number of k-mer scores calculated were evaluated to find the best-performing combination. The optimal parameters were (1) to use the longest ORFs with either a start codon, a stop codon or both (ORFs of type 3, following the notation of the FEELnc publication); and (2) to calculate multi k-mer scores of k-mers of k in {1, 2, 3, 6, 9, 12}, instead of individual ones. Finally, and like CPAT, a 10-fold cross-validation was used to calculate the optimal coding probability cutoff, which is automatically used by the tool to label transcripts. FEELnc was evaluated on the test set together with other classification tools, producing a sensitivity of 0.923, specificity of 0.915 and an F1-score of 0.919. It is important to note that, instead of providing a trained model, the FEELnc tool trains a RF classifier on the go using the provided training set.

We used FEELnc version 0.2.1 to train and evaluate a RF model on our dataset. We ran FEELnc following the optimal parameters provided in the publication, including the use of 'type 3' ORFs and the calculation of multi k-mer scores. The probability cutoff found was 0.4202.

##### *LncDC*

LncDC is a gradient boosting classifier based on the XGBoost library [19]. This model leverages a set of sequence-intrinsic, secondary structure and protein features. Interestingly, LncDC predicts ORFs in 4 distinct ways (called ORF types) and calculates sequence and protein features for all of them. The LncDC training and testing datasets were constructed as follows: Protein coding transcripts were collected from RefSeq release 99 and lncRNAs from Gencode v34. Transcripts were filtered out by three criteria: (1) length < 200 nt, due to the definition of lncRNAs as transcripts longer than 200 nt (2) length > 20000 nt, due to the exponential computational cost of SS calculation and (3) presence of nucleotides other than 'ACGT'. Then, a held-out test set was constructed with 10000 protein-coding transcripts and 10000 lncRNAs. The rest of the sequences were scanned with CD-HIT to remove those with an identity higher than 90%, limiting sequence redundancy. The remaining 37,698 mRNAs and 34,539 lncRNAs were used to build the training dataset. In addition, to balance the number of transcripts in both classes, a set of synthetic lncRNA sequences was created using the SMOTE technique, which recursively creates sequences by selecting random pairs of instances in the feature space and choosing a random point between them.

A set of 57 features was evaluated by training an XGBoost model and performing random feature elimination with cross-validation (RFECV). A subset of 28 features was found to have the maximum accuracy (0.9823) while minimizing the number of features, avoiding potential feature redundancy and overfitting. This final set contained (1) sequence features, such as the GC-content, the Fickett score, and metrics of different ORF types (maximum ORF length, ORF coverage, hexamer scores and relative codon bias); (2) secondary structure features, including the GC-content of paired nucleotides and the sequence and secondary-structure (SASS) k-mer scores for k values in [1,5], and protein features, like the isoelectric point, molecular weight, aromaticity and instability of the translations of different ORF types.

The final model was trained and evaluated with the data sets described above. LncDC achieved a sensitivity score of 0.9812, specificity of 0.9786 and an F1-score of 0.9799.

We trained a LncDC model using version 1.3.6 with our custom dataset. Since the tool allows for training either with or without SS features, we built those models and evaluated their performance on our held-out data set. Like in the LncDC publication, the model integrating SS features provides a marginal performance improvement, so we continued working with this version (Table S6). Importantly, model training produces custom files that are key for later model inference, including an hexamer table, an imputer for over-sampling of the minority class and a scaler to standardize feature values as z-scores using training data feature distribution.

Note that we included minor modifications in the source code of LncDC to allow it to load precalculated secondary structure sequences, reducing training and inference time. These modifications can be found in our fork of the LncDC code (https://github.com/cbib/LncDC, commit ID 8cbce76)

##### *Lncfinder*

Lncfinder is a toolbox to evaluate transcript features and train transcript classifiers [17]. It distributes a trained support vector machine (SVM) classifier, hereby called Lncfinder. Lncfinder incorporates a total of 19 features, including (1) sequence-intrinsic features (the logarithmic distances (LDs) of hexamer composition, the length and the coverage of ORFs), (2) secondary structure features (MFE, frequency on unpaired nucleotides followed by a paired nucleotide or UP frequency, and the LDs of different encodings of secondary structure); and (3) physicochemical features based on the fast fourier transform (FFT) of electron-ion interaction pseudo-potential (EIIP) (the signal at ⅓ position, the signal to noise ratio and the signal quantile statistics min, Q1, Q2 and max). Key features within each group were selected after RFECV or a custom selection algorithm to later validate the global performance of combining all 19 features.

The LncFinder publication uses datasets described elsewhere (Achawanantakun et al., 2015). Briefly, Human dataset A (or H1 in the original publication) contains 15308 protein-coding transcripts and 4586 lncRNAs in the training set, all extracted randomly from single genes from the Gencode database. The testing set is made of 4000 transcripts of each class, with none of them sharing genes with training transcripts. Additionally, Human dataset B (or H2) is a modified version of the CPAT dataset (see above), with a training set reduced to 9929 mRNAs and 9066 lncRNAs due to database updates, and an equivalent testing set. Feature selection and architecture comparison was performed by 10-fold CV using the training set of Human dataset A, and final model training and benchmarking used the training and testing sets of Human dataset B, respectively. Evaluation of LncFinder SVM resulted in a sensitivity of 0.9544, specificity of 0.9150 and F1-score of 0.9360.

We trained a custom SVM using LncFinder version 1.1.6. Due to the computational constraints of the RNAfold tool used for SS calculation, sequences longer than 30000 nucleotides were excluded for the analysis (Table S17). Given that the SS predicted by RNAfold may not be the actual molecular structure and to include the long transcripts, we evaluated the performance of the a LncFinder model without SS information. The metrics were slightly reduced, so we kept the SS model for further experiments (Table S6).

##### *PlncPRO*

PlncPRO is a transcript classification tool, originally developed for plants, based on random forests[10]. PlncPRO is trained on a set of 71 features, including:

- 64 trimer frequencies

- 4 metrics extracted from BLASTX hits against SWISS-PROT. These were (1) number of hits (N); (2) significance score (S) calculated as the sum of negative natural logarithm of the e-value for each BLASTX hit; (3) the sum of bitscores of all his (B); and (4) frame entropy (F), calculated as the Shannon entropy of the probability that a hit falls within a given reading frame.
- 2 ORF-based features from Framefinder, including the ORF coverage and the Framefinder score (FF-score).

The PlncPRO program trains 5 independent RF classifiers and keeps the one with highest out-of-bag score, a metric that estimates generalization error. Finally, the selected model is used for transcript classification. The PlncPRO publication describes the training and evaluation of multiple models trained on transcripts of a range of plant species. Moreover, they report the performance of a similar model trained and tested on Human transcripts. This model achieves accuracies of 0.9434 and 0.9372 on two Human test sets.

Model retraining was performed as per the PlncPRO instructions. However, due to an unidentified bug, BLASTX features were not properly computed for the training set, which may negatively impact classification performance.

##### *mRNN*

mRNA RNN (mRNN) is a modified recurrent neural network (RNN) model that uses gated recurrent units (GRU) to learn sequential information from transcripts and classify them [12]. mRNN encodes sequences as embedding vectors and uses dropout regularization to reduce overfitting. This model was trained and evaluated on a series of datasets created from Gencode v25 transcripts: (1) a test set with 500 mRNAs and 500 lncRNAs longer than 200nt extracted from individual genes, (2) a “challenge” set consisting of mRNAs with ORFs shorter than 150 nt and lncRNAs longer than 200 nt containing a non-coding ORFs of at least 150 nt, (3) a “long 5’ UTR set” comprising the 500 mRNAs with longest 5’UTRs; from the remaining sequences, (4) a validation set with 500 coding and 500 non-coding transcripts, (5) a training set only including sequences between 200 and 1000 nt, with a total of 16000 transcripts of each class; and (6) an augmented pre-training dataset constructed by introducing 10 mutated copies of training transcripts, each with one random 1-nt insertion. Pre-training was performed on sets 6 and 4, model training was done with datasets 5 and 4, and sets 1, 2 and 3 were used in final model evaluation. It is important to note that pre-training set augmentation by random insertions and restriction of training sequence lengths were reported to provide improvements in classification performance. The training process entailed (1) pre-training 30 models with distinct augmented datasets for 4 epochs, (2) evaluating pretrained models on the validation dataset to select the top 6, (3) further train the best models on the non-augmented dataset for 10 epochs during 10 independent sessions and (4) select the top 5 final models, keeping only one resulting model per pre-trained dataset. Finally, transcript classification is performed either by using the best single model (mRNN) or integrating the top 5 models by averaging model weights (mRNN ensemble). The ensemble model achieved higher scores in all metrics analysed, including specificity, sensitivity and F1-score higher than 0.95 (average of 100.000 bootstrap trials with replacement).

We modified the published version of the mRNN code to increase the maximum allowed transcript length, which was hardcoded and caused an infinite loop if longer sequences were used. Code modifications can be found in our fork (https://github.com/cbib/mRNN). We used this modified version of mRNN (commit 41fc34a) to train an ensemble classifier. The common training dataset was further split into pre-training and validation datasets, with the latter containing the same number of transcripts as the testing set. In order to keep an equivalent training set for all tools evaluated in this study, the training set was not filtered by transcript size. Then, data augmentation, model pre-training, pre-trained model selection, model training and final model evaluation were performed following the same protocol as the original publication.

##### *RNAsamba*

RNAsamba is a deep learning model that incorporates the IGLOO architecture [18]. Its main innovation is the combination of two independent branches: B1 branch, that uses the nucleotide sequence (trimmed to 3,000 nucleotides) to capture coding signals independent of ORF identification; and the B2 branch, that integrates the amino acid sequence encoded by the longest ORF (truncated at 1,000 aa) together with sequence features, including nucleotide k-mer frequencies, amino acid relative frequencies and ORF length. These two branches are combined through a learned attention parameter that dynamically weights their relative contribution, which is particularly important for transcripts where no ORF was identified.

The RNAsamba authors published two trained models: one trained on full transcript sequences, and a second one trained on both full-length and truncated transcripts. Train/test sets were extracted from previous publications, including CPC2, FEELnc, mRNN and CPPred. Sequence redundancy was limited by running MMseq with 90% homology and 90% coverage filters. Model evaluation was performed against CPC2, CPAT, FEELnc, lncRNAnet and mRNN on several datasets, and RNAsamba scored the highest area under the precision-recall curve (AUC) for all of them, with AUC values between 94.16% and 99.67%.

We trained an RNAsamba model using full transcripts following the original instructions. The choice to use full sequences instead of truncated transcripts was motivated by the fact that our benchmark is based on reference datasets instead of on experimental transcript models that would increase fragmentation.

##### *LncRNA-BERT*

lncRNA-BERT is a BERT-based RNA language built on the transformer encoder architecture with a binary classification head [23]. It supports supports two sequence encoding strategies:

- K-mer tokenization (k=3): overlapping trinucleotide tokens from the raw sequence (64-token vocabulary).
- Convolutional Sequence Encoding (CSE) (k=9): a learnable convolutional layer that embeds raw nucleotide sequences into a continuous token space, providing stronger compression that allows longer transcripts to be classified.

Since transformer architecture is limited by the context size, lncRNA-BERT trims the sequence end to make it fit.

The model training is composed of two steps:

- Pre-training (masked language modeling): The model learns general RNA sequence representations from GENCODE v46, RefSeq v255, and NONCODE v6 (a total of 297,724 coding and 238,470 non-coding sequences). 5% of GENCODE is held out for validation.
- Fine-tuning (binary classification task): The pre-trained model is fine-tuned for coding/non-coding classification on 101,270 coding and 48,785 lncRNA sequences from GENCODE and RefSeq (NONCODE excluded to maximise data reliability). Importantly, sequence redundancy in the fine-tuning set was limited by using CD-HIT with a 90% identity threshold.

lncRNA-BERT models were evaluated on three testing datasets: (1) GENCODE/RefSeq set, (2) CPAT test set and (3) the multi-species RNAChallenge dataset of transcripts difficult to classify.

Both k-mer tokenization and CSE encoding versions were evaluated, with the following respective F1 scores: (1) 0.940 and 0.943, (2) 0.963 and 0.947; and (3) 0.235 and 0.242. Importantly, while the performance for the RNAChallenge dataset is significantly lower to the other datasets, lncRNA-BERT models achieved the best performance over all models compared in their study (i.e., CPAT, LncFinder, PredLnc, LncADeep, RNAsamba and mRNN).

For our benchmark, we used the pre-trained model with the k-mer tokenization (k=3) and fine-tuned each validation fold with its corresponding training dataset. Since the fine-tuning step requires a validation dataset, similarly to mRNN, we held-out a subset of transcripts from the training set for this purpose.

#### *Supplemental figure*

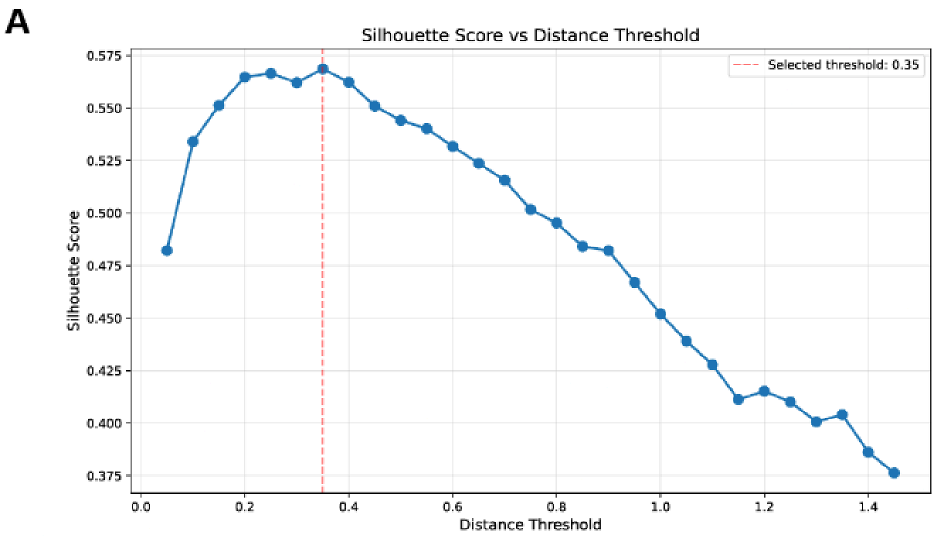

**Figure S2. Silhouette score of feature clusters.** (A) Silhouette score was calculated for incremental distance thresholds, and the highest was retained as optimal to define the number of clusters (155 clusters at a distance of 0.35)

#### *Supplemental tables*

| **Table S5:** Number of features after each processing step | | | | |
|---|---|---|---|---|
| Step | ML Set | Rep Set | NBD Set | Total |
| Raw tables | 172 | 173 | 189 | 534 |

| Table S5: Number of features after each processing step | | | | |
|---|---|---|---|---|
| **Numerical features** | **128** | **170** | **178** | **476** |
| Non-constant | 128 | 157 | 178 | 459 |
| Continuous | 128 | 142 | 169 | 435 |
| Representative | 66 | 36 | 53 | 155 |
| Categorical | 0 | 15 | 9 | 24 |
| **Final set** | **66** | **51** | **62** | **179** |

**Table S17:** Transcripts in the “common CDHIT” set without missing classification labels

| tool | count | reason |
|---|---|---|
| total | 112,264 | (All transcripts in the common cdhit dataset) |
| total no class | 612 | (All transcripts that were not classified by all tools) |
| ss_lncDC | 529 | Sequence shorter than 200 nt, containing non-ACGT characters or longer than 20,000 nt. |
| l_cpat | 180 | ORF not found or sequence shorter than 75 nt. |
| feelnc | 5 | ORF not found |
| ss_lncfinder | 15 | Sequence longer than 30,000 nt. |

Tables S3, S4 and S6 to S16 are available in the Supplemental Tables file.

### Categorization of transcripts based on confidence and agreement

***Supplemental figure***

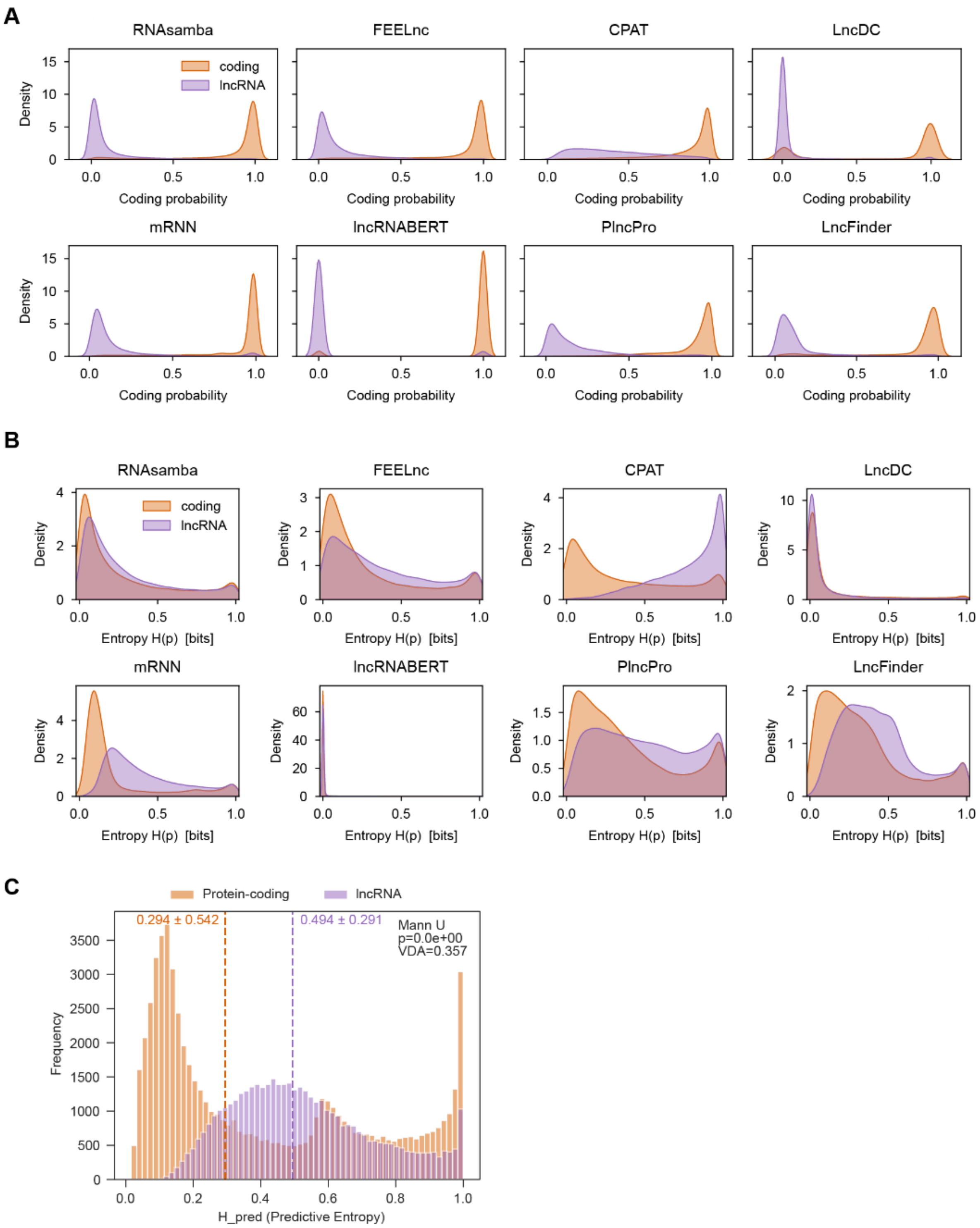

**Figure S3. Model probabilities and entropy distributions.** (A) Distribution of model probabilities, where 1 = protein-coding and 0 = lncRNA. (B) Distribution of model entropy, where 0 = full certainty and 1 = full uncertainty. (C) Distribution of predictive entropy ($H_{pred}$).

### Incorporation of repetitive elements information

#### *Supplemental methods*

RepeatMasker v4.2.1, with its default human database (Dfam version 3.9) as reference, was used to annotate the repetitive elements present in the transcript. RepeatMasker was run with the following parameters: species set to “human”, sensitive mode (-s) and parallel processing. The Gencode FASTA headers were trimmed to the first pipe character (“|”) to ensure compatibility.

RepeatMasker output files (“.out”) containing hits were parsed. To keep a single hit annotation per coordinate, overlapping hits were resolved following a hierarchical scoring system. RM reports some elements that overlap with a higher-scoring hit with an asterisk in the last column of the “.out” file. For overlapping pairs, we removed (1) hits tagged with an asterisk by RM; if neither had an asterisk (2) hits with lower Smith-Waterman alignment score (SW score); if scores were equal, (3) hits with lower percentage overlap. If overlaps remained, we resolved them by keeping the first reported element.

Then, using a derivative of the Dfam classification, hits were separated into four groups detailed in the following table:

**Table S18:** Groups of repetitive elements used in the feature extraction pipeline

| Group | Class |
|---|---|
| Transposable Elements | DNA, LINE, LTR, PLE, RC, Retroposon, SINE and srpRNA* |
| Interspersed repeats and low complexity regions | Satellite, Simple_repeat and Low_complexity |
| Pseudogenes | rRNA, scRNA, snRNA and tRNA |
| Unknown | Unknown |

* Due to the absence of clear classification of srpRNAs in the classification tree of Dfam v3.9

| **Table S18:** Groups of repetitive elements used in the feature extraction pipeline |
| --- |
| (https://www.dfam.org/releases/Dfam_3.9/infrastructure/TEClasses.tsv), we included srpRNAs as a standalone class within the Transposable Elements class. This choice was informed by the fact that all srpRNA elements are 7SLRNA in Dfam family DF000000016, which is included in the Alu class. |

Then, a comprehensive set of 87 features was extracted using a custom Python pipeline. Feature extraction proceeded hierarchically at three levels:

Hit-level features: individual RepeatMasker hits were characterized by SW score, divergence from the consensus sequence, insertions and deletions (all RM features). We further calculated hit length and reference coverage (proportion of the consensus repeat covered by the hit).

Element-level features: RepeatMasker reports fragmented hits that represent nested insertions or incomplete elements. Fragments belonging to the same original element were aggregated using RepeatMasker's hit identifiers. For each reconstructed element, we computed total length, total reference coverage, number of fragments and fragmentation status (a binary indicator of having >1 fragment). Element-level classifications were performed for seven major TE classes: LINE (Long Interspersed Nuclear Elements), SINE (Short Interspersed Nuclear Elements), LTR (Long Terminal Repeats), DNA transposons, RC (rolling-circle transposons), Retroposons and PLE (Penelope-like elements). Element classifications included family resolution, capturing Alu, MIR, L1, L2 and ERV subfamilies among others. Elements were additionally classified as "young" (L1HS, L1PA1–2, AluY, SVA_F, HERVK) or "ancient" (divergence >20%).

Transcript-level features: All elements overlapping each transcript were aggregated to generate transcript-level summary statistics. For each transcript, we computed: (1) count-based features including total number of hits, number of fragmented elements, fragmentation ratio and diversity metrics (number of unique subfamilies, families and classes); (2) length-based features including total TE coverage (bp and percentage), and minimum/mean/maximum (min/mean/max) values for hit lengths and reference coverage values; (3) divergence-based features including min/mean/max values for SW scores, sequence divergences, deletions and insertions; (4) class-presence features including hit counts and presence/absence binary indicators for each major class and family; and (5) spatial features including gap statistics (min/mean/max and standard deviation of inter-element distances) and hit density (hits per kilobase).

Additionally, we calculated features relative to transcript length (hit length, min/mean/max hit length, and gap stats) and per kilobase (total hit count, class/family hit count and unique class/family count). All 158 extracted features are comprehensively detailed in Table S3.

### Incorporation of non-B DNA structure information

#### *Supplemental methods*

Non-B DNA structural motifs were identified on the GRCh38 genome reference assembly using two tools.The *non-B_gfa* (gfa) pipeline (https://github.com/abcsFrederick/non-B_gfa), which generates annotation files for for A-phased repeats (APR), direct repeats (DR), inverted repeats (IR), mirror repeats (MR), short tandem repeats (STR), and Z-DNA. Triplex-forming motifs (TRI) were defined as the subset of MR predictions flagged as triplex-prone by the gfa output. G-quadruplexes (G4) were predicted on GRCh38 using G4Discovery [24] with default parameters, reporting both pqsfinder [25] and G4Hunter [26] scores.

All motif calls were intersected with GENCODE v47 transcript intervals (genomic coordinates) using bedtools intersect (v7.3.0), retaining all pairwise overlaps (-wa -wb). The resulting files were fed into a feature extraction pipeline that (1) clips NBD motifs extending beyond transcript genomic boundaries, (2) resolves overlapping elements of the same motif class, (3) computes a panel of features for each motif class and (4) computes cross-motif gap statistics. Features per motif included:

- Presence indicator
- Hit count
- Min, max, mean and standard deviation of motif length.
- Total motif length, both before and after overlap resolution.
- Min, max, median, mean and standard deviation of inter-motif gaps
- Relative features as percentage of the transcript length and counts per kilobase

The full set of 178 extracted features is comprehensively detailed in Table S4.
